# Context-dependent role of vinculin in neutrophil adhesion, motility and trafficking

**DOI:** 10.1101/847780

**Authors:** Zachary S. Wilson, Hadley Witt, Lauren Hazlett, Michael Harman, Brittany M. Neumann, Andrew Whitman, Mohak Patel, Robert S. Ross, Christian Franck, Jonathan S. Reichner, Craig T. Lefort

## Abstract

Neutrophils are innate immune effector cells that traffic from the peripheral blood to extravascular sites of inflammation. β2 integrins are involved during multiple phases of neutrophil recruitment, including the transition from rolling to arrest, firm attachment and motility within the vasculature. Following neutrophil arrest, adhesion stabilization occurs as the neutrophil interacts with the endothelial surface and crawls into a favorable position for extravasation. The cytoskeletal protein vinculin has been implicated in other cell types as a regulator of adhesion strength by promoting focal adhesion maturation and as a sensor of the mechanical properties of the microenvironment. Neutrophils express vinculin but do not form mature focal adhesions. Here, we characterize the role of vinculin in β2 integrin-dependent neutrophil adhesion, motility, mechanosensing, and recruitment. We observe that knockout of vinculin attenuates, but does not completely abrogate, neutrophil adhesion, spreading, and crawling under static conditions. In the presence of forces from fluid flow, vinculin was not required for neutrophil adhesion or migration. Vinculin deficiency only mildly attenuated neutrophil traction stresses and spreading on stiff, but not soft, polyacrylamide gels indicating a minor role for vinculin in the mechanosensing of the neutrophil as compared to slower moving mesenchymal cells that form mature focal adhesions. Consistent with these findings, we observe *in vivo* neutrophil recruitment into the inflamed peritoneum of mice remains intact in the absence of vinculin. Together, these data suggest that while vinculin regulates some aspects of neutrophil adhesion and spreading, it may be dispensable for neutrophil recruitment and motility *in vivo*.

## Introduction

Neutrophils are leukocytes of the innate immune system that are the first to respond and mobilize to sites of infection or injury. The recruitment of neutrophils from the circulation is mediated by β2 integrins that interact with endothelium adjacent to an inflamed tissue site (1, 2). Humans that lack β2 integrins or their activators suffer from leukocyte adhesion deficiency type I and III, respectively, which greatly increases host susceptibility to bacterial and fungal opportunistic pathogens (1–3). However, excess recruitment and retention of neutrophils at sites of inflammation can also lead to bystander injury through the release of reactive oxygen species and proteolytic enzymes from preformed granules (4, 5). Many investigators have studied the mechanisms of neutrophil migration to find therapeutic targets that might enable tighter control over the inflammatory response without impairing host defense. With this long-term goal in mind, this study probes the role of vinculin in neutrophil adhesion, motility, and trafficking mediated by β2 integrins.

In the classical model of neutrophil recruitment, expression of P- and/or E-selectin on inflamed endothelium mediates the initial tethering and rolling of neutrophils. During rolling, neutrophils receive activation signals via engagement of P-selectin glycoprotein ligand-1 (PSGL-1) and G protein-coupled receptors, such as the canonical neutrophil chemokine receptors for IL-8 (human) or CXCL1 (murine). In a process called “inside-out” activation, these signals trigger structural changes in β2 integrins that increase their ligand-binding affinity by up to four orders of magnitude (6, 7). High affinity β2 integrins, primarily LFA-1 (CD11a/CD18), mediate the transition of rolling neutrophils to arrest and firm adhesion by binding to intercellular adhesion molecule-1 (ICAM-1) (8). Finally, arrested neutrophils will then spread and crawl toward a favorable site for transmigration (9). In this way, β2 integrin inside-out signaling, where intrinsic signals through PSGL-1 and CXCR2 activate β2 integrins, dominate the steps leading to neutrophil arrest, while β2 integrin “outside-in” signaling downstream of ICAM-1 engagement, are critical for stabilizing adhesion, intraluminal spreading and crawling (10).

Vinculin is a scaffolding protein involved in the maturation of integrin-based focal adhesions that has been studied primarily in mesenchymal cells such as fibroblasts. Vinculin has multiple binding surfaces to enable the recruitment of proteins to adhesion sties (11). Vinculin is maintained in an inactive state through intramolecular association of its head and tail domains; activation through intramolecular dissociation is thought to be mediated by phosphorylation and sequential binding to a series of proteins (12–15). The localization of vinculin at integrin-mediated adhesions can occur through its association with phosphorylated paxillin that shuttles vinculin to talin-1 that is concomitantly bound to the tail of an activated integrin (16). Vinculin stabilizes integrin adhesions within a mature focal adhesion through the recruitment of actin-binding proteins and the direct binding of actin bundles (17). Vinculin-dependent focal adhesion maturation has been described as being “mechanosensitive,” which refers to the recruitment of vinculin through actomyosin-mediated contractility and transmission of signals that scale with the mechanical stiffness of the substrate (18). The current study establishes a mechanosensitive role for vinculin during neutrophil adhesion and spreading mediated by β2 integrins.

Although vinculin function has been studied in other leukocytes, this is the first study to investigate the potential role for vinculin in leukocyte trafficking (19–22). Activated neutrophils express vinculin and form focal complexes, but they are also highly motile amoeboid-like cells that do not generate mature focal adhesions (23, 24). Adhesion stabilization enables neutrophils to crawl toward favorable sites of emigration, but whether this process is vinculin-dependent is unclear (25). We report that while vinculin contributes to neutrophil adhesion to ICAM-1 in the absence of shear stress, it is dispensable for neutrophil adhesion and motility under shear stress, and for infiltration into the inflamed peritoneum of mice. In addition, we show that, as in other cell types, vinculin plays a role rigidity sensing for neutrophils, as measured by their spread area and traction force generation. Together, these data point towards a less prominent role for vinculin in neutrophils, as compared to mesenchymal cells, that depends on the properties of the microenvironment.

## Materials and Methods

### Antibodies and reagents

All antibodies used are against murine antigens. Antibodies: anti-CD18 (clone GAME-46; BD Biosciences), anti-CD11a (clone M17/4; BioLegend), anti-ICAM-1 (clone YN1; BioLegend), APC-anti-CD11b (clone M1/70; BioLegend), anti-Ly6G (clone 1A8; BioLegend), APC-anti-CD117 (clone 2B8; BioLegend), anti-CXCR2 (clone SA045E1; BioLegend), anti-α-actinin (Cell Signaling Technologies), anti-vinculin (Cell Signaling Technologies), anti-GFP (Cell Signaling Technologies), HRP-conjugated-anti-Rabbit IgG (Cell Signaling Technologies), anti-CD11a (clone IBL-6/2; Cell Signaling Techologies), Alexa Fluor 647-anti-Rat IgG (ThermoFisher Scientific). Reagents: recombinant murine CXCL1 (BioLegend), recombinant murine SCF (BioLegend), recombinant murine G-CSF (BioLegend), recombinant murine ICAM-1 (R&D Systems, BioLegend), 4-Hydroxytamoxifen (Tocris), CFSE (BioLegend), TagIt-Violet (BioLegend), Thioglycollate broth (Sigma Aldrich), PKH26/PKH67/Claret Far Red Membrane Dye (Sigma Aldrich).

### Neutrophil progenitors

Neutrophils were obtained by differentiating murine myeloid progenitors that were conditionally-immortalized using tamoxifen-inducible HoxB8 (26). Briefly, murine hematopoietic stem/progenitor cells were isolated from bone marrow (StemCell Technologies), transduced with a tamoxifen-inducible expression vector for the murine *Hoxb8* gene (27), and then cultured in the presence of 100 nM 4-Hydroxytamoxifen (4-OHT), 50 ng/mL recombinant murine stem cell factor, and 1 μg/mL puromycin (26). Progenitors were differentiated into neutrophils by removing 4-OHT and culturing in the presence of 20 ng/mL recombinant murine stem cell factor and 20 ng/mL recombinant murine granulocyte colony-stimulating factor for 4 days (Fig. S1A) (28). Neutrophils differentiated from progenitors exhibit multi-lobed nuclei, expression of Ly6G, and a loss in the expression of CD117 (cKit) (Fig. S1B-C). To create a vinculin knockout progenitor cell lines for this study, HoxB8-conditional progenitors were transduced with a lentiviral vector that expresses Cas9 and single-guide RNA (sgRNA) targeting the *Vcl* gene that encodes vinculin. To do so, we used the pLentiCRISPR v2 vector, a gift from Feng Zhang (Addgene plasmid #52961) that was modified to confer blasticidin resistance, and the following sgRNA target sequences: TTCCCCTAGAGCCGTCAATG (Vcl (1)) and CCGGCGCGCTCACCCGGACG (Vcl (2)) (Fig. S2A). The *Tln1* and *Itgb2* genes were knocked out in HoxB8-conditional progenitors as previously described (29). Empty vector expression of Cas9 without a targeting sgRNA was used as a wild-type control for all experiments. Vinculin was successfully disrupted in progenitors after a single lentiviral transduction that was followed by blasticidin selection (Fig. S2B). When using a fluorescent reporter of HoxB8 expression, no difference was observed between wild-type (WT) and vinculin knockout (VclKO) EGFP-HoxB8 expression before differentiation and both exhibited a similar loss of EGFP-HoxB8 expression at the end of four days of differentiation (Fig. S2C).

For rescue studies, Clover (a GFP variant) conjugated to vinculin or the vinculin A50I mutant were cloned into the doxycycline-inducible Tet-On 3G plasmid system (Takara Bio). HoxB8-conditional progenitors (expressing Cas9 and control sgRNA or Vcl (2) sgRNA) were transduced by lentivirus with the inducible Clover-vinculin constructs and then treated with doxycycline (Sigma Aldrich) at a concentration of 1 μg/mL to induce expression of Clover-vinculin. After transduction, progenitors were sorted for high expression of Clover-vinculin using fluorescence activated cell sorting (FACS). Clover-vinculin was successfully expressed in both control and VclKO progenitors derived from Vcl (2) sgRNA that targets the intron-exon junction, and therefore does not target exogenous Clover-vinculin (Fig. S3).

### Neutrophil static adhesion assay

Neutrophils obtained after 4-day differentiation (“*in vitro*-derived”) were washed and labeled using CFSE (BioLegend). Murine bone marrow neutrophils were isolated by negative selection (StemCell Technologies) and immediately labeled alongside *in vitro*-derived neutrophils using CFSE. 96-well plates were coated for 1 hour at room temperature or overnight at 4°C with 2.5, 5, or 7.5 µg/mL ICAM-1 and/or 2.5 µg/mL CXCL1 in phosphate buffered saline (PBS) and then blocked with 1% casein (ThermoFisher) or 0.5% polyvinylpyrrolidone (Sigma Aldrich) in PBS for 2 hours at room temperature. Neutrophils were loaded into the 96-well plate at 0.5×10^6^ neutrophils per well in Hank’s balanced salt solution containing Ca^2+^ and Mg^2+^ (HBSS^++^), and then incubated at 37°C for 35 or 65 minutes. Following incubation, neutrophils were quantified by a plate reader for fluorescence intensity (CFSE), before and after sequential gentle washes with HBSS^++^. The number of adherent neutrophils was inspected visually by light microscopy to corroborate with plate reader signal intensity. Each group was replicated in three to six wells per independent experiment.

### Neutrophil motility and spreading on glass

Neutrophils obtained after 4-day differentiation were washed and labeled using CFSE (BioLegend). Delta T dishes (Bioptech) were coated with 2.5 µg/mL ICAM-1 and 2.5 µg/mL CXCL1 in PBS overnight at 4°C and blocked for 1 hour with 0.5% PVP in PBS at room temperature. Approximately 75,000 neutrophils were added to warm HBSS^++^ and migration was followed using time-lapse microscopy for 30 minutes at 37°C with images captured every 20 seconds. Motility was tracked using ImageJ Manual Tracking and analyzed using Ibidi Chemotaxis to obtain measures of migration such as accumulated distance, Euclidean distance, and velocity. Cell area was measured using the final image acquired at 30 minutes using ImageJ.

### Neutrophil spreading on polyacrylamide gels

Polyacrylamide gel substrates were prepared as originally described (30). Briefly, gels were prepared using varying concentrations of acrylamide (Bio-Rad) and *N,N*-methylene-bisacrylamide (Bio-Rad) to achieve the range of elasticity. Stiff substrates were chosen to be 12% acrylamide and 0.4% bisacrylamide, at approximately 100 kPa stiffness; intermediate stiffness substrates were chosen to be 8%/0.08%, at approximately 8.3 kPa; and soft substrates were chosen to be 3%/0.2%, at approximately 1.5 kPa. Within the smaller range of intermediate stiffness gels, acrylamide/bisacrylamide used were: 6.4%/0.23% for approximately 20 kPa stiffness, 5.2%/0.19% for approximately 10 kPa, and 4.4%/0.16% for approximately 5 kPa. Polyacrylamide solutions were vortexed and then polymerized through the addition of excess tetramethylethylenediamine and ammonium persulfate. Gels were polymerized at room temperature on hydrophilic-treated, glass-bottom deltaT dishes (Bioptech) and shaped using AbGene frames to a final size of approximately 1 cm × 1 cm × 300 µm. Gels were then soaked for at least one hour in water to remove any unpolymerized acrylamide. Final elasticity was measured by atomic force microscopy. Sulfo-SANPAH was allowed to covalently bond to the gel for 1 hour in the dark. ICAM-1 and/or CXCL1 were then UV cross-linked to the sulfo-SANPAH at 2.5 µg/mL each. Previous work shows that final elasticity is not affected by surface protein crosslinking, and that final protein density is not affected by elasticity (31). After washing with buffered saline, *in vitro*-derived neutrophils were added in warmed HBSS^++^ and incubated at 37°C. Imaging was performed 30 minutes after incubation. For motility assays, imaging was performed within a fully enclosed microscope at 37°C for 30 minutes in L-15 media supplemented with 2 mg/ml glucose.

### Flow chamber assay

To prepare flow chambers, Ibidi µ-Slide VI^0.1^ were coated with 0.5 μg/mL E-selectin, 7 μg/mL ICAM-1, and 8 μg/mL CXCL1 for 2 hours in PBS and then blocked with an excess of casein for 2 hours, both at room temperature. Flow chambers were perfused at 12.98 µL/min, which is calculated to produce a shear stress of 1 dyne/cm^2^. CFSE-labeled wild-type and vehicle control-treated vinculin knockout neutrophils were evaluated within the same flow chamber. Time-lapse images were captured every 10 seconds for 1 hour, starting immediately after starting flow chamber perfusion, using transmitted light through a 20X objective. Motility was tracked using ImageJ Manual Tracking and analyzed using Ibidi Chemotaxis to obtain measures of migration: accumulated distance, Euclidean distance, and velocity.

### Western blot

For each group, 1×10^6^ neutrophils were lysed in Tris lysis buffer (0.2M Tris, 1% Triton X-100, 1% sodium orthovandate) containing protease inhibitor cocktail (Sigma Aldrich) for 30 minutes on ice with occasional mixing. Lysed samples were centrifuged at 11,000g for 10 minutes and the supernatants were heated to 95°C for 10 minutes in Laemmli buffer. Supernatants were loaded onto 4-20% Mini-PROTEAN TGX gels (Bio-Rad) and run at 160V. Gels were then transferred onto 0.45 µm nitrocellulose membrane at 100V for 1 hour at 4°C. Probing of western blots was carried out according to the antibody manufacturer’s instructions (Cell Signaling Technologies). Western blots were analyzed with SuperSignal West Pico PLUS Chemiluminescent Substrate (Thermo Scientific) to detect HRP-conjugated antibodies using a Bio-Rad Chemidoc XRS.

### Immunocytochemistry and TIRF microscopy

Glass coverslips (0.17 mm) were coated with 10 µg/mL ICAM-1 and 2.5 µg/mL CXCL1 for 2 hours and then blocked with an excess of casein for 2 hours. Neutrophils obtained after 4-day differentiation were washed and resuspended in HBSS^++^ prior to use. Neutrophils were incubated on coverslips for 35 minutes at 37°C, then fixed in 10% neutral buffered formalin for 30 minutes, and permeabilized in 0.1% Triton X-100 for 10 minutes. Coverslips were incubated with primary antibodies in 1% BSA overnight at 4°C, transferred to secondary antibodies for 1 hour at room temperature. Neutrophils were stained with NucBlue and ActinGreen 488 (ThermoFisher Scientific) according to manufacturer’s recommendation prior to mounting coverslips onto slides. For live cell imaging, progenitors with transduced to express Clover-vinculin and Lifeact-mRuby2 in differentiated neutrophils. Samples were imaged with a TILL Photonics iMIC TIRF microscope (FEI Company) and Andor iXon3 EMCCD camera.

### Animals

All animal studies were approved by the Lifespan Animal Welfare Committee. Mice were housed in a specific pathogen-free facility at Rhode Island Hospital. Mice harboring floxed *Vcl* alleles (Vcl^f/f^) were kindly provided by Dr. Robert Ross (UC-San Diego) and have been previously described (32). Vcl^f/f^ mice were crossed with MX1-Cre (MX1^cre^) mice (The Jackson Laboratory) in which Cre recombinase expression is controlled by the MX1 promoter and can be induced by interferon production after administration of synthetic double-stranded RNA (33). To generate mixed chimeras, 8- to 12-week-old C57BL/6 mice (The Jackson Laboratory) were lethally irradiated (10 Gy, single dose) and then reconstituted by intravenous injection of bone marrow cells from a Vcl^f/f^MX1^cre^GFP^+^ mouse expressing transgenic enhanced green fluorescent protein (EGFP) under the ubiquitin C promoter (The Jackson Laboratory) and a Vcl^f/f^ (GFP^-^) control mouse at 1:1 ratio. Deletion of the gene encoding vinculin was induced by intraperitoneal injection of 250 μg of polyinosinic–polycytidylic acid (Poly I:C; InvivoGen), three doses, each 2 days apart, starting 4 weeks after irradiation, inducing near complete loss of the respective protein in neutrophils (Fig. S4). Mice were used for experiments 4–8 weeks after Poly I:C administration.

### Peritonitis model

For each mixed chimeric mouse, a blood sample was collected by saphenous vein puncture prior to intraperitoneal injection of 1 mL 4% thioglycollate broth to induce peritonitis. For adoptive transfer studies, mice were first challenged with an intraperitoneal dose of 1 mL 4% thioglycollate broth. At 2 hours post-challenge, mice were intravenously injected with a mixed 1:1 population of 6 × 10^6^ membrane dye-labeled *in vitro*-derived neutrophils. At the indicated time point, 5 mL of ice-cold PBS with 2 mM EDTA was used to lavage the peritoneum. Blood and lavage were analyzed by flow cytometry using fluorescently labeled anti-Ly6G antibody to distinguish neutrophils with a MACSQuant Analyzer 10 (Miltenyi).

### Cremaster muscle intravital imaging

Mice were anesthetized using a cocktail of ketamine (125 mg/kg) and xylazine (12.5 mg/kg), the carotid artery was cannulated, and the cremaster muscle was exteriorized, cut longitudinally, and spread onto a stage as has been previously described (7). The cremaster muscle was perfused throughout the experiment with 37°C bicarbonate buffered saline equilibrated with 5% CO_2_ in N_2_. A blood sample was collected prior to imaging using a catheter placed in the carotid artery. For arrest assays, 600 ng murine CXCL1 was intravenously administered through the catheter. Cremaster muscle post-capillary venules were imaged for 14 minutes following CXCL1 injection using an upright Olympus BX60 microscope with a 40X water-immersion objective with EGFP fluorescence captured by a Chameleon 3 color camera (FLIR Systems) and Rapp SP20-X3 xenon flashlamp.

### Traction force microscopy

Traction force microscopy was performed as previously described, with some modifications (34). Briefly, neutrophils obtained after 4-day differentiation were washed, labeled using CFSE (ThermoFisher Scientific), and resuspended in HBSS^++^. Polyacrylamide gels were prepared using varying concentrations of acrylamide (Bio-Rad) and *N,N*-methylene-bisacrylamide (Bio-Rad) to achieve the range of desired elasticities with a Poisson’s ratio of 0.5. Soft gel substrates were composed of 3% acrylamide and 0.2% bisacrylamide at approximately 1.5kPa, very soft substrates were composed of 3% acrylamide and 0.06% bisacrylamide at approximately 0.5 kPa. Gel substrates were embedded with 0.2% final w/v 0.5 µm fluorescent microspheres (Invitrogen). Polyacrylamide gels were crosslinked using 1.25% APS and 0.5% TEMED yielding a final gel thickness of approximately 40 µm. 100 µg/mL ICAM-1 and 5 µg/mL CXCL1 were UV cross-linked to the sulfo-SANPAH, as described above. Three-dimensional time-lapse volumetric images of fluorescent beads in polyacrylamide substrates and labeled cell membranes were recorded using laser scanning confocal microscopy (LSCM). A fast iterative digital volume correlation (FIDVC) algorithm was used to track the motion of beads and calculate the 3D displacement vector during cell migration compared to a reference position obtained by detaching the cells using 2% sodium dodecyl sulfate (Sigma Aldrich) (34).

To calculate cell tractions, **T**, and an overall measure of cellular contractility, *μ*, the Cauchy stress, **σ**, was determined using the following equation for a Neo-Hookean elastic solid,

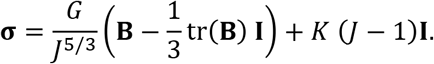

Here, *J* is the Jacobian of the deformation gradient tensor, **B** is the left Cauchy-Green deformation tensor, and *G* and *K* are the shear and bulk modulus of the polyacrylamide gel, respectively. To compute the surface normal, **n**, we applied the Delaunay triangulation on the deformed surface grid points. The reference gel surface is assumed to be initially flat in the stress-free-state, and we compute the deformed surface grid points by adding the cumulative surface displacement to the reference gel surface.

Using the surface normal, **n**, and the Cauchy stress, **σ**, we compute the cell tractions, **T**, and root-mean squared traction, *T_rms_*, as follows,

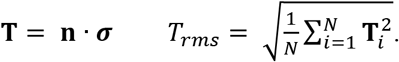

To establish a simple, scalar-based metric of the overall contractility of the cell, we extended the general procedure by Butler et al. to calculate the three-dimensional dipole moment tensor as,

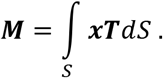

The cellular contractility, *µ*, is then defined simply as the trace of the dipole moment tensor **M**, i.e., *µ* = tr(**M**) (35).

### Data analysis

All analyses were performed using GraphPad Prism 8. As indicated, one-way or two-way analysis of variance (ANOVA) was used to compare the differences between samples and post-hoc analysis was performed using Tukey pairwise multiple comparison test. For samples that did not have a normal distribution, Kruskal-Wallis one-way ANOVA was used to compare difference between samples and post-hoc analysis was performed using Tukey pairwise multiple comparison test. Experimental data are presented with mean and standard deviation. For traction force microscopy, µ is log transformed as this better describes the distribution of values.

## Results

Neutrophils are terminally differentiated leukocytes with a limited lifespan and therefore not amenable to genetic manipulation for *in vitro* studies. To circumvent this limitation, we first established the utility of HoxB8-conditional myeloid progenitors as an *in vitro* source of differentiated mature neutrophils, progenitor-derived wild-type (WT) and vinculin knockout (VclKO) neutrophils were examined for markers to confirm their neutrophil identity and function. Ly6G is a murine neutrophil-specific marker and its expression level corresponds with bone marrow differentiation of precursors into mature neutrophils (36). For all HoxB8-conditional progenitor cell lines used in this study, differentiation in the presence of SCF and G-CSF resulted in the complete loss in expression of CD117 and the gain in the expression of Ly6G (Fig. S1A). LFA-1 and Mac-1, the two primary β2 integrins expressed by neutrophils, consist of the common β2 subunit CD18 and α subunits CD11a and CD11b, respectively. We observed similar expression of CD11a and CD11b by progenitor-derived WT, VclKO (1) and (2) neutrophils created using two distinct sgRNAs, and by bone marrow (BM) neutrophils isolated from mice (Fig. S5A-B). Surface expression of CD11a and CD11b was ablated in β2 integrin knockout (Itgb2KO) neutrophils, as previously characterized (29). All groups of progenitor-derived neutrophils expressed the canonical chemokine receptor CXCR2 (Fig. S5C). In examining the activation of neutrophils, the upregulation of CD11b from delivery of intracellular granule stores to the cell surface was found to be similar in both WT and VclKO neutrophils, with an approximate 4-fold increase in expression in response to formylated peptide Met-Leu-Phe (fMLP) (Fig. S5D) (37). Altogether, these data establish the *in vitro*-derivation of genetically-modified murine neutrophils from conditionally-immortalized progenitors.

To determine whether vinculin plays a role in β2 integrin-mediated adhesion, a static adhesion assay was used to measure neutrophil attachment to a substrate of ICAM-1 and CXCL1. We observed that vinculin-deficient neutrophils had attenuated adhesion compared to WT neutrophils, and had comparable levels of adhesion as neutrophils lacking β2 integrin expression (Fig. 1A). Adhesion levels of progenitor-derived WT neutrophils were not statistically different from that of BM neutrophils (Fig. 1A). The reduction in adhesion of vinculin-deficient neutrophils was observed over a range of substrate ligand concentrations and assay wash stringency (Fig. S6A-C). Overall, these results suggest that vinculin plays a role in β2 integrin-mediated adhesion by neutrophils.

**Figure 1.**
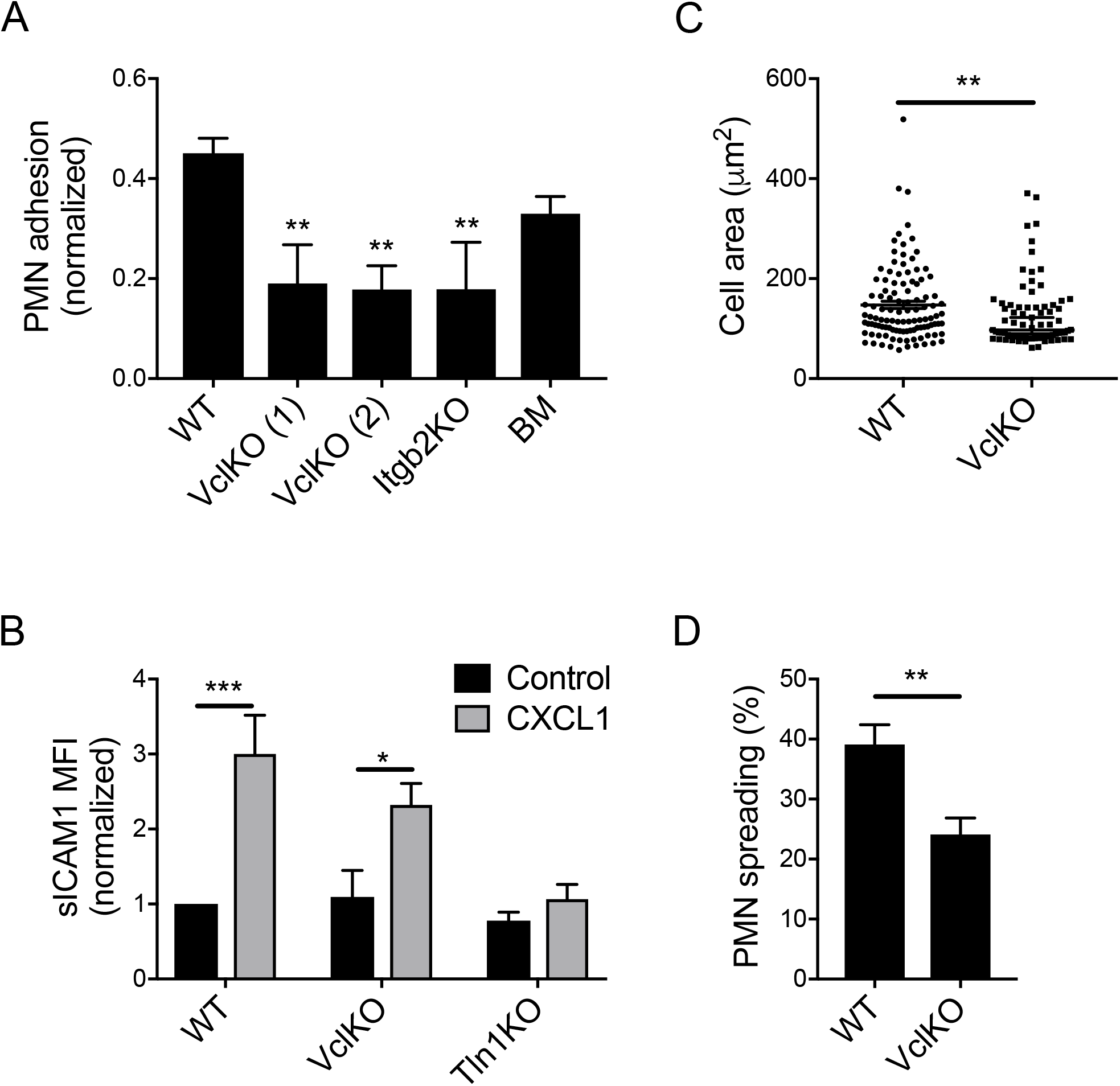
Vinculin knockout attenuates β2 integrin-dependent neutrophil adhesion. (A) Adhesion of neutrophils to immobilized ICAM-1 and CXCL1 assessed for progenitor-derived wild-type (WT), vinculin knockout (VclKO) created using sgRNAs (1) and (2), β2 integrin knockout (Itgb2KO), and murine bone marrow neutrophils (n=3 independent experiments). Analyzed using one-way ANOVA with Tukey pairwise multiple comparison test. ** p<0.01. (B) Soluble ICAM-1 binding to neutrophils in response to CXCL1, as measured by flow cytometry (3 replicates per group, n=3 independent experiments). Analyzed using two-way ANOVA with Tukey pairwise multiple comparison test. * p<0.05; *** p<0.001. (C) Spread area of membrane-labeled neutrophils on immobilized ICAM-1 and CXCL1 (n>70 cells/group, n=2 independent experiments). Analyzed using Mann-Whitney Rank Sum Test. ** p<0.01. (D) Percent of neutrophils spreading on immobilized ICAM-1 and CXCL1 after 30 minute incubation (n>17 fields of view/group, n=3 independent experiments). Analyzed using unpaired Student’s t-test. ** p<0.01.

On the surface of quiescent neutrophils in the circulation, β2 integrins are in an inactive state with low ligand-binding affinity. In response to activation signals received during selectin-mediated rolling on inflamed endothelium, “inside-out” activation of β2 integrins triggers a drastic increase in binding to ICAM-1 that mediates the transition from neutrophil rolling to arrest (8). As an assay specific for the detection of β2 integrin affinity changes, soluble ICAM-1 binding has been previously used to demonstrate the essential role of talin-1 and Kindlin-3 in this process (7). As VclKO neutrophils had impaired adhesion relative to WT, we sought to determine whether vinculin plays a role in the inside-out activation of β2 integrins by measuring soluble ICAM-1 binding in response to CXCL1. As expected, WT neutrophils exhibited a significant increase in soluble ICAM-1 binding in the presence of CXCL1, while talin-1-deficient neutrophils were unable to activate their β2 integrins to bind soluble ICAM-1 (Fig. 1B). Vinculin-deficient neutrophils responded to CXCL1 in a similar manner as WT neutrophils and bound significantly more ICAM-1 than unstimulated neutrophils, indicating that β2 integrin activation remains intact in the absence of vinculin (Fig. 1B). Levels of CXCL1-induced soluble ICAM-1 binding by VclKO neutrophils were not significantly different from WT neutrophils (Fig. 1B). These data indicate that while vinculin regulates neutrophil adhesion, it is not involved in the earliest steps of adhesion that rely on inside-out β2 integrin activation.

Neutrophil spreading occurs prior to migration where the neutrophil flattens along its substrate and increases its integrin-mediated associations (38). Spreading involves activated integrin clustering and actin rearrangements (39, 40). Vinculin plays a role in actin polymerization, integrin clustering, and adhesion plaque maturation in other cell types (17, 41). To understand whether vinculin plays a similar role in neutrophil spreading, we quantified cell area and frequency of spreading under the same conditions as those used in evaluating neutrophil adhesion. We observed that spreading is impaired in VclKO neutrophils, in terms of both cell area and the fraction of cells that spread beyond the diameter of a round neutrophil in suspension (Fig. 1C-D). Thus, vinculin is required for efficient and complete neutrophil spreading on ICAM-1 in response to CXCL1.

Migration of neutrophils is dependent on both adhesion turnover and actin polymerization (42, 43). Vinculin has been previously described to regulate directed motility in fibroblasts (44). Here, we examined neutrophil migration to understand the potential role of vinculin in neutrophil motility during chemokinesis. As compared to WT neutrophils, VclKO neutrophils exhibited mild attenuation of migration with significantly lower accumulated distance, Euclidean distance, and instantaneous velocity (Fig. 2A-C). Directness of migration, a measure of the tendency of the neutrophil to travel in a straight line, was similar for wild-type and vinculin-deficient neutrophils (Fig. 2D). These data suggest that neutrophils generally undergo random-walk chemokinetic behavior under these conditions, as directness below 0.5 indicates less than half of migration is in the direction of its ending position. Individual neutrophil migration tracks are shown in Figure 2F. To better analyze this behavior, a two-dimensional algorithm for measuring single particle diffusion was used to calculate mean-squared displacements for migration up to 80 seconds. Mean-squared displacement of VclKO neutrophils was significantly impaired compared to WT for migration up to 80 seconds (Fig. 2E). After 80 seconds the error for this model in both wild-type and vinculin knockout neutrophils is inflated and therefore unreliable. Altogether, these data indicate a role for vinculin in neutrophil migration under static conditions.

**Figure 2.**
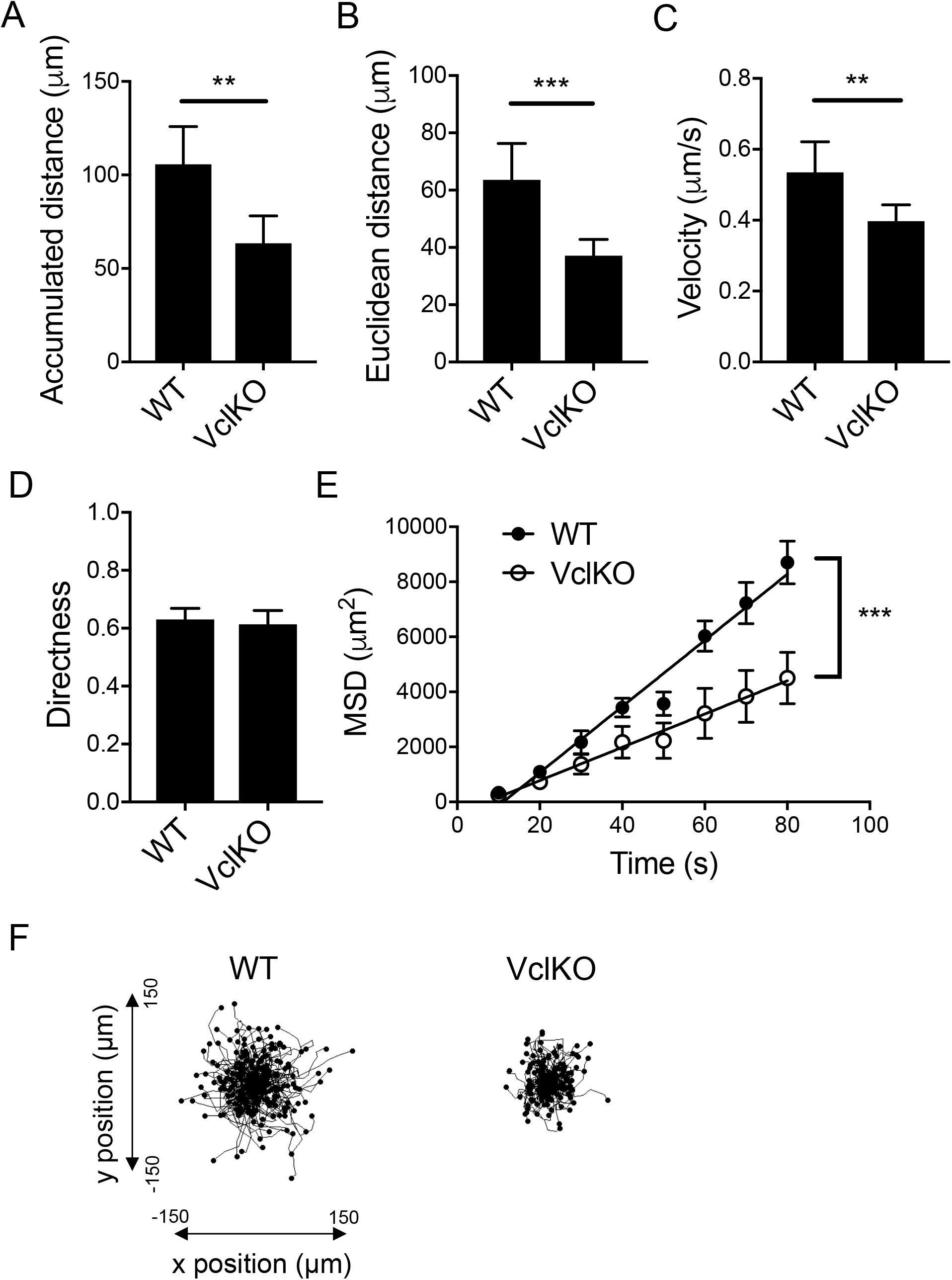
Vinculin plays a role in β2 integrin-dependent neutrophil motility. (A-D) Parameters of neutrophil motility during 30-minute chemokinesis on immobilized ICAM-1 and CXCL1 (n>160 cells/group, 3 independent experiments). Analyzed using unpaired Student’s t-test. ** p<0.01; *** p<0.001. (E) Mean-squared displacement of neutrophil migration based on particle modeling. Analyzed using linear regression with comparison of slopes. WT: y = 120t – 1338; VclKO: y = 60.5t - 431.3. *** p<0.001. (F) Individual neutrophil tracks during 30-minute chemokinesis on immobilized ICAM-1 and CXCL1.

In the context of neutrophil recruitment from the circulation, migration on ICAM-1 expressed on the endothelium will typically occur in an environment in which forces from blood flow are experienced by the attached neutrophil. Thus, to better recapitulate physiological conditions, we performed neutrophil migration assays in a flow chamber perfused at a wall shear stress within the range typical of post-capillary venules. Flow chambers were coated with E-selectin, ICAM-1, and CXCL1 to reconstitute ligands presented by inflamed endothelial cells that mediate neutrophil rolling, arrest, and intraluminal migration. In contrast to the apparent defect in VclKO neutrophil migration under static conditions, no significant difference was observed in the accumulated distance, Euclidean distance, and instantaneous velocity of WT and VclKO neutrophils in the presence of fluid shear stress (Fig. 3A-C and Supplementary Video 1). WT and VclKO neutrophil directness were similar (Fig. 3D), and the Rayleigh p-value was below 0.001 for both WT and VclKO neutrophils, suggesting that both groups of neutrophils were similarly moving in the direction of fluid flow rather than in random chemokinetic motion (Figure 3E). Thus, in the presence of forces from fluid flow, vinculin plays no apparent role in β2 integrin-mediated neutrophil motility.

**Figure 3.**
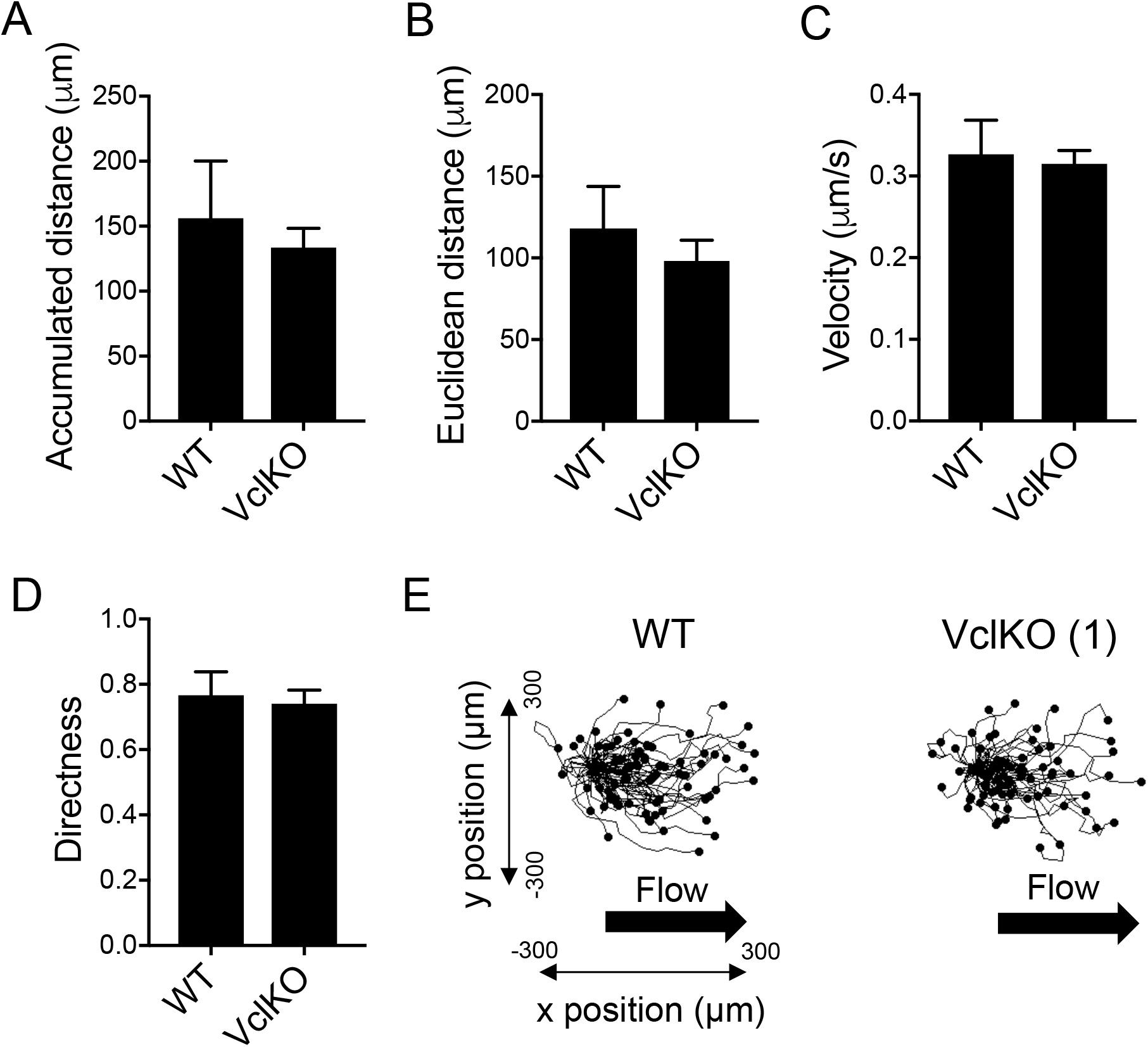
Vinculin is dispensable for neutrophil motility under shear stress. (A-D) Parameters of neutrophil motility during 60-minute chemokinesis in a flow chamber coated with E-selectin, ICAM-1 and CXCL1, and perfused at a wall shear stress of 2 dyne/cm^2^ (n>70 cells/group, 5 replicate runs, 2 independent experiments). Analyzed using unpaired Student’s t-test. (E) Neutrophil tracks during 60-minute chemokinesis in a flow chamber (as in A-D).

To understand whether the impairment in spreading and migration are related to the actin cytoskeleton, WT and VclKO neutrophils adherent to immobilized ICAM-1/CXCL1 were examined for the localization of actin by fluorescence microscopy. CD11a, the α subunit of LFA-1, was used to examine integrin localization (25). Phalloidin was used to stain F-actin, which is expected to localize to uropods as stable F-actin fibers and as actively polymerizing in the lamellipodia during migration (43, 45). Total internal reflection fluorescence microscopy (TIRFM) was used to selectively image fluorescent signals at the cell surface interacting with the substrate. WT neutrophils had a strong localization of F-actin within uropods, where force is expected to be generated for ameboid migration (46), while there was no discernable organization pattern of CD11a (Fig. 4A). In observing actin distribution, 77% of WT neutrophils could be considered polarized compared to 32% and 25% in VclKO (1) and (2), respectively (Fig. 4B and S7A). The fluorescent skewness and kurtosis of F-actin distribution was measured to determine whether the distribution of fluorescence was asymmetric around the mean or peaked, respectively. Skewness of greater than 1 can be considered asymmetrical while positive kurtosis indicates peaked intensity away from a Gaussian distribution. F-actin median skewness in WT neutrophils was observed to be 1.23 compared to VclKO (1) and (2), which have a median skewness of 0.89 and 1.02, respectively (Fig. S7B). All neutrophils displayed similar median kurtosis values (Fig. S7B). After normalizing to WT, VclKO (1) and (2) neutrophils had a median F-actin intensity in TIRFM images that was significantly less than that of WT neutrophils (Fig. 4C), whereas CD11a intensity was not different across groups (Fig. 4D). Neutrophil morphology was measured based on aspect ratio, roundness and circularity; higher values of aspect ratio and lower values of roundness and circularity imply more elongated neutrophil morphology. The median circularity, median roundness and median aspect ratio of WT neutrophils were significantly different than that of VclKO neutrophils (Fig. S7B). Altogether these data indicate that vinculin-deficient neutrophils are less polarized compared to WT neutrophils, which may be related to impairment in actin cytoskeletal organization. When vinculin was expressed endogenously with a fluorescent tag or visualized using antibody labeling, it was found to localize to the perimeter of the neutrophil (Fig. S8A-B), as has been previously described (24, 47). Using neutrophils expressing Clover-vinculin and Lifeact-mRuby2, to visualize actively polymerizing actin, surface-proximal vinculin was tracked in a live neutrophil migrating on ICAM-1/CXCL1 (Fig. S8A). We observed that Clover-vinculin increased in intensity as neutrophils contracted inward during migration, suggesting a potential role for vinculin during the contraction stage of migration.

**Figure 4.**
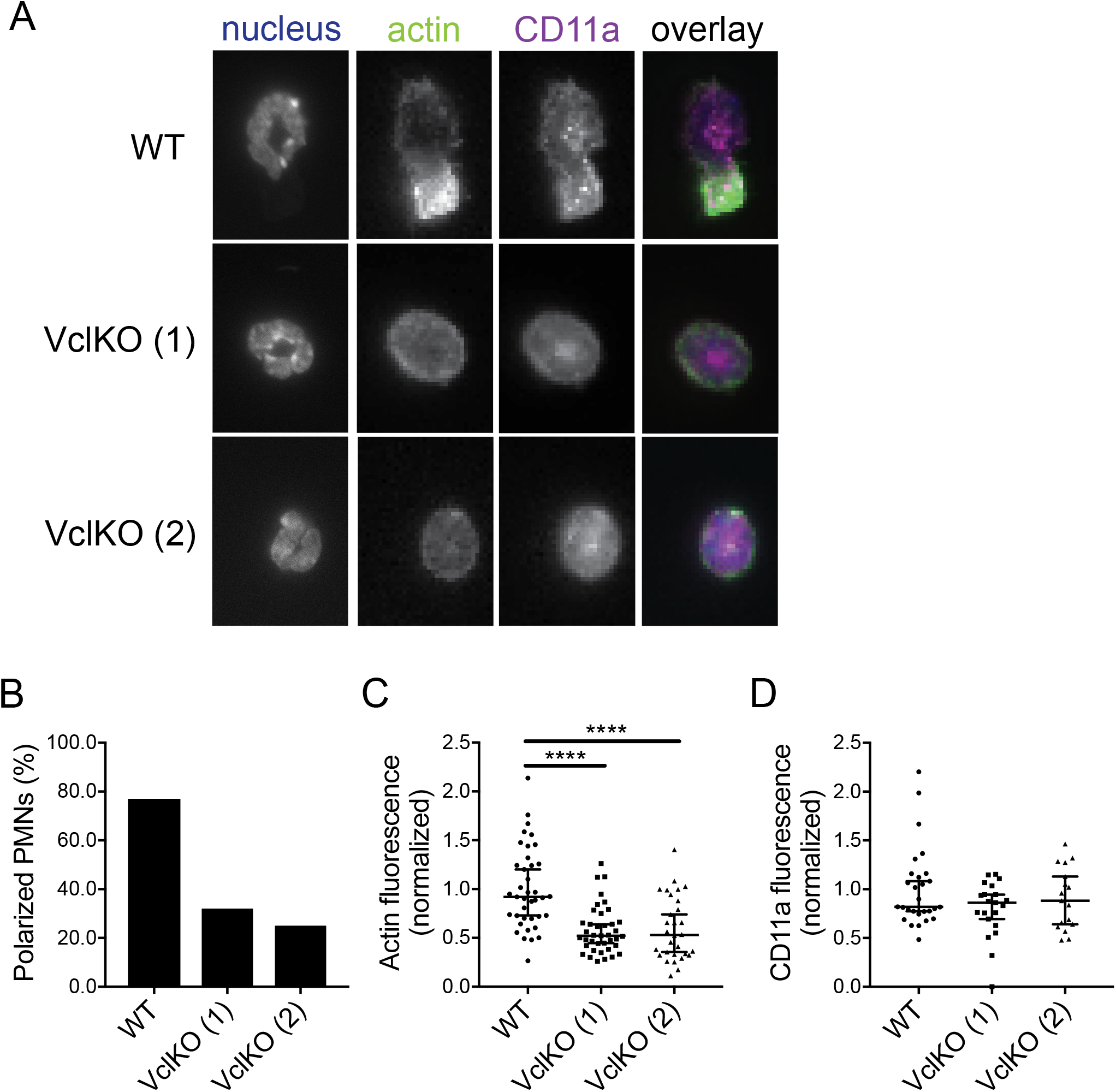
Vinculin is required for stable F-actin localization within uropods and neutrophil polarization. (A) Representative immunoflluorescence images of neutrophils after 30-minute incubation on immobilized ICAM-1 and CXCL1. Cells were stained with Hoescht, phalloidin, and anti-CD11a (n=3 independent experiments). Scale bar = 10 µm. (B) Polarization of neutrophils based on asymmetric F-actin distribution (n=3 independent experiments). (C-D) Background-subtracted and normalized TIRFM fluorescent intensities of F-actin and CD11a (n>30 cells/group, 3 independent experiments). Analyzed using Kruskal-Wallis one-way ANOVA with Dunn’s multiple comparison test. **** p<0.0001.

To better understand the apparent contradictory results on the role of vinculin in neutrophil migration, a murine mixed bone marrow chimeric model was used to examine whether vinculin regulates *in vivo* neutrophil recruitment. Mixed chimeric mice allow for analysis of competitive *in vivo* recruitment of wild-type (Vcl^f/f^) and vinculin knockout (Vcl^f/f^MX1^cre^) neutrophils in an internally controlled inflammatory environment. For mice challenged with an intraperitoneal injection of thioglycollate broth that induces sterile inflammation, neutrophil recruitment peaks during the first 4-6 hours and occurs through mechanisms involving β2 integrins (1). Comparing the baseline (pre-stimulus) peripheral blood chimerism to that observed in the peritoneal lavage at 4 hours after inducing peritonitis, we found no significant difference in the recruitment of wild-type and vinculin-deficient neutrophils (Fig. 5A). Additionally, *in vitro* progenitor-derived WT and VclKO neutrophils were adoptively transferred into C57BL/6 mice to observe their competitive recruitment during thioglycollate-induced peritonitis. Again, there was no difference in the recruitment of *in vitro*-derived WT and VclKO neutrophils, whereas Itgb2KO neutrophils exhibited impaired recruitment into the inflamed peritoneum as expected (Fig. S9C-D).

**Figure 5.**
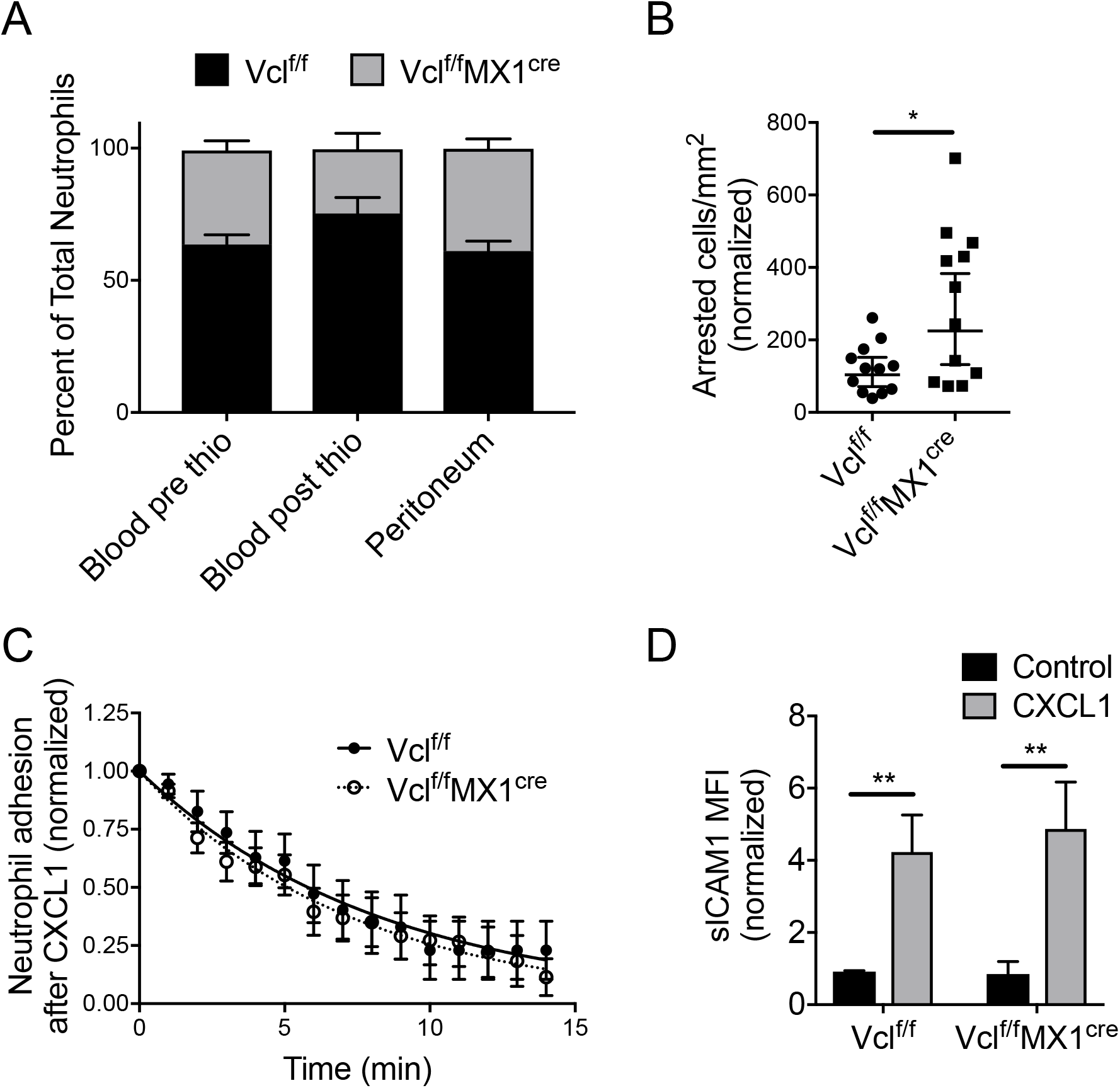
Vinculin is dispensable for neutrophil recruitment *in vivo*. (A) Percentage composition of control (Vcl^f/f^) and vinculin knockout (Vcl^f/f^MX1^cre^) neutrophils in the peripheral blood and peritoneal lavage, 4 hours after induction of peritonitis in mixed chimeric mice (n=5 mice, 2 independent experiments). Analyzed using two-way ANOVA with Tukey multiple comparison test. (B-C) The arrest and time course of sustained adhesion of neutrophils in response to intravenous injection of CXCL1, and over the 15-minute period immediately following (n=12 fields of view, across 7 chimeric mice). Data were analyzed using non-linear regression: (WT) Y = e^-0.119t^, (VclKO) Y = e^-0.136t^. (D) Soluble ICAM-1 binding to bone marrow neutrophils in response to CXCL1, as measured by flow cytometry (3 replicates per group, n=3 independent experiments).

The murine cremaster muscle microvasculature was observed in mixed chimeric mice by intravital microscopy during soluble chemokine (CXCL1) stimulation, which has previously been shown to induce rapid β2 integrin-mediated neutrophil arrest that is dependent on integrin activation by talin-1 and Kindlin-3 (7). In addition, the time that elapses prior to detachment of neutrophils after CXCL1-indued arrest was used to measure adhesion strengthening (48). We observed no impairment in the arrest or adhesion strengthening of vinculin-deficient neutrophils relative to wild-type (Fig. 5B-C). Further, Vcl^f/f^ and Vcl^f/f^MX1^cre^ bone marrow neutrophils assayed *ex vivo* exhibited a similar increase in soluble ICAM-1 binding (Fig. 5D), supporting previous results using progenitor-derived neutrophils and indicating that vinculin is not required for β2 integrin activation. It is therefore unclear how vinculin deficiency results in enhanced numbers of neutrophils arresting on post-capillary venules after CXCL1 stimulation (Fig. 5B). Altogether, these *in vivo* data are consistent with our *in vitro* findings analyzing neutrophil migration under fluid flow that suggest vinculin is dispensable for intraluminal neutrophil motility. Further, these *in vivo* data also suggest that the process of neutrophil diapedesis and entry into extravascular tissue sites does not require vinculin.

Vinculin is well characterized for its mechanosensitive function in other cell types, and so we reasoned that vinculin may play an analogous role in neutrophils (49, 50). The spreading of human neutrophils has been shown to depend on substrate stiffness (31), but the molecules involved in the neutrophil mechanosensing response have yet to be identified. To probe the function of vinculin in neutrophil mechanosensing, we analyzed neutrophil spreading on polyacrylamide gels of varying stiffness that were functionalized with ICAM-1 and CXCL1. With increasing substrate stiffness, WT neutrophils exhibited an increase in cell area and the fraction of neutrophils that spread, whereas VclKO neutrophils exhibited an attenuated response that differed significantly from WT neutrophils only at the highest (100 kPa) substrate stiffness (Fig. 6A-B and S10A). While the mechanosensitive spreading of VclKO neutrophils was significantly attenuated relative to WT, vinculin deficiency did not completely ablate β2 integrin-dependent spreading, as demonstrated by comparing VclKO neutrophils to Itgb2KO neutrophils lacking β2 integrin expression (Fig. 6A-B). We further probed neutrophil adhesion and motility at intermediate substrate compliance (5 – 20 kPa), observing that the mechanosensitive phenotype of vinculin-deficient neutrophils became measurable in our experimental system within this intermediate range of substrate stiffness (Fig. S10B). In addition, we observed that neutrophils lacking vinculin had impaired motility on 5 kPa and 10 kPa gels, but not on 20 kPa gels (Fig. S10C).

**Figure 6.**
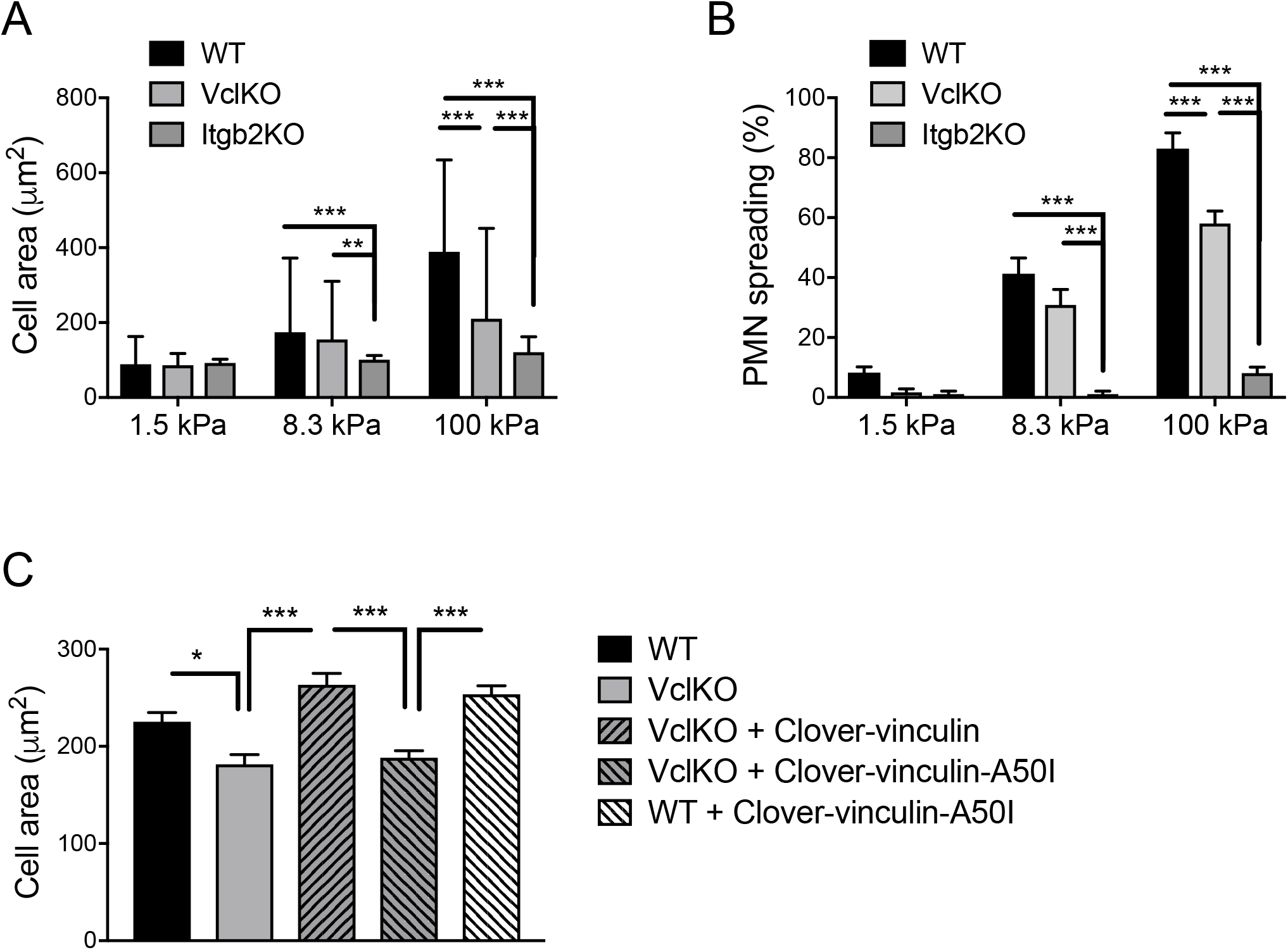
Vinculin plays a role in neutrophil mechanosensing. (A-B) Neutrophil cell area and spreading frequency on polyacrylamide gels of varying stiffness (soft: 1.5 kPa, intermediate: 8.3 kPa, stiff: 100 kPa) conjugated with ICAM-1 and CXCL1 (n>100 cells/group, 15 fields of view/group, 3 independent experiments). Analyzed using two-way ANOVA with Tukey multiple comparison test. ** p<0.01, *** p<0.001, **** p<0.0001. (C) Area of neutrophils from the indicated groups of WT and VclKO (2) cells with exogenous expression of Clover-vinculin in wild-type or A50I mutant forms. Neutrophils were allowed to adhere and spread on 100kPa polyacrylamide gels conjugated with ICAM-1 and CXCL1 (n>250 cells/group, 12 fields of view/group, 3 independent experiments) * p<0.05, *** p<0.001.

To gain further insight into the mechanisms of vinculin-mediated neutrophil mechanosensing and spreading, we attempted rescue of VclKO neutrophil spreading on 100 kPa substrates by exogenous expression of wild-type or mutant forms of vinculin. Expression of vinculin tagged with a variant of GFP, Clover-vinculin, was able to enhance the spread area of VclKO neutrophils to levels observed in WT neutrophils (Fig. 6C). However, expression of Clover-vinculin-A50I, with a single amino acid mutation that disrupts interaction with talin-1 (17), was not able to rescue the spreading deficiency of VclKO neutrophils (Fig. 6C). Altogether, these data indicate that vinculin regulates β2 integrin-dependent neutrophil spreading through a mechanosensing mechanism that involves vinculin interaction with the integrin tail-binding protein talin-1.

Contractile force generation is essential for neutrophil adhesion, spreading, and migration (42). To quantify contractility, we performed traction force microscopy using bead-embedded polyacrylamide gels. There was a technical limitation for these studies, in that only on gel substrates of relative low stiffness (less than 1.5 kPa) do neutrophils produce measurable gel/bead displacements. Nevertheless, traction force microscopy is a sensitive technique capable of resolving small differences in traction stress that do not necessarily manifest in a population-level phenotype. Polyacrylamide gels were functionalized as above with ICAM-1 and CXCL1, but with 40-fold more ICAM-1 and 2-fold more CXCL1 to maximize contractility in each individual cell. Possibly due to this increased ligand density, neutrophils underwent adhesion and spreading, but there was no observable long-range migration under any of the measured conditions. *In vitro*-derived WT and VclKO neutrophils generated increased traction stresses from very soft (0.5 kPa) to soft (1.5 kPa) gels, but had similar overall contractility under the tested conditions (Fig. 7A-B). VclKO neutrophils had reduced traction stresses on soft gels compared to WT neutrophils, but similar overall contractility for both very soft and soft gels. These data are consistent with VclKO neutrophil spreading being unimpaired on gels of lower matrix relative to WT, suggesting again that the magnitude of the role for vinculin in neutrophil adhesive function depends on the mechanical microenvironment.

**Figure 7.**
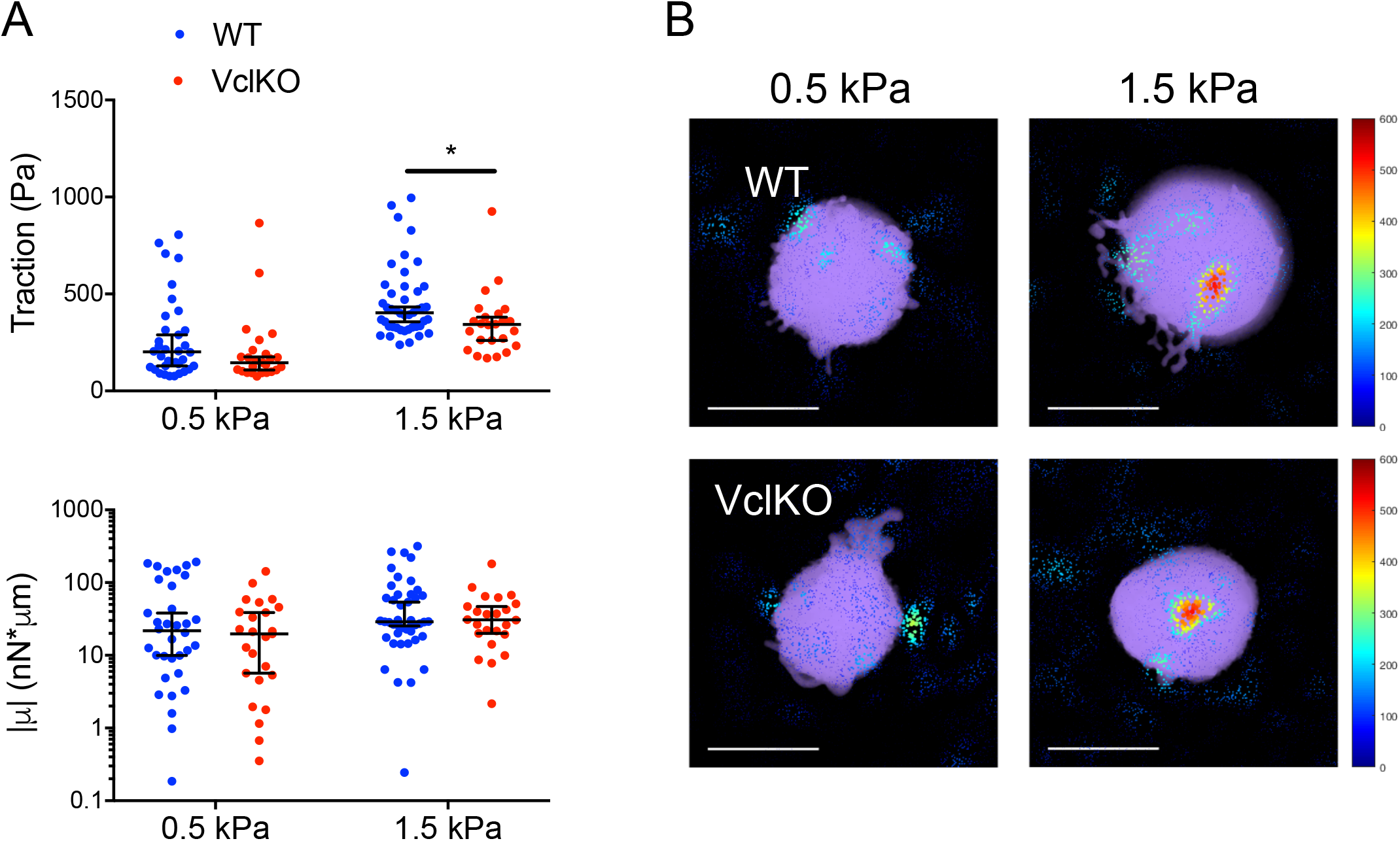
Neutrophil traction stress generation is attenuated by vinculin deficiency. (A) 3D tractions and the trace of the dipole moments, µ, of WT and VclKO neutrophils on polyacrylamide gels of either 0.5 kPa or 1.5 kPa stiffness (n>25 cells/group, 3 independent experiments). Data analyzed using two-way ANOVA with Tukey multiple comparison test. *p<0.05. (B) Representative 3D traction cone plots for WT and VclKO neutrophils (shown in purple) on polyacrylamide gels of indicated stiffness. Cones indicate traction direction while color and size represent traction magnitude, in Pascals. Scale bar =10 µm.

## Discussion

The goal of this study was to assess the role of vinculin in neutrophil adhesion and motility. In the classic example mesenchymal cell adhesion, vinculin is involved in the maturation of integrin-based focal adhesions and contributes to cell spreading and mechanotransduction (51, 52). In leukocytes, vinculin has been shown to have roles in unique processes like T and B cell immune synapse formation, marking apoptosis for T cells, and osteoclast actin-ring formation during bone resorption (20, 22, 53). In platelets, vinculin is dispensable for most physiological functions (54). Whether vinculin is a mediator or regulator of neutrophil adhesion was uncertain prior to the current study. Although neutrophils do not form mature focal adhesions, we hypothesized that vinculin may play a role in neutrophil behavior during processes such as adhesion strengthening to the endothelium.

By live cell imaging, we observed that vinculin accumulates in punctate structures in the neutrophil as it contracts inward on ICAM-1, similar to what has been previously described for vinculin localization in neutrophils (55). Our study finds that vinculin knockout attenuates neutrophil adhesion, spreading and migration on ICAM-1 *in vitro* under static conditions. The attenuated response in adhesion mimics the attenuation in spreading, which may imply that the neutrophils remaining adhered in the static adhesion assay are those that are spread. Despite the *in vitro* defect observed in neutrophils lacking vinculin, there was no *in vivo* phenotype as assessed in a classical recruitment model of acute peritonitis. Furthermore, neutrophil adhesion strengthening in inflamed post-capillary venules of the cremaster muscle remained intact in vinculin-deficient neutrophils. In fact, there were significantly more vinculin knockout neutrophils that underwent rapid arrest in post-capillary venules in response to CXCL1, an *in vivo* assay of β2 integrin activation. It is possible that in mixed chimeric mice the supply of vinculin knockout neutrophils to the cremaster muscle vascular bed was transiently enhanced relative to wild-type, as neutrophils can become sequestered elsewhere in the circulation in response to systemic stimuli.

During sterile inflammation, chemokines form a gradient outward from the offended site that guides migrating neutrophils (56). Neutrophils first encounter these chemokines intravascularly as they are immobilized by heparan sulfate on the apical surface of the inflamed endothelium (56). Neutrophil crawling on inflamed endothelium is necessary to find a favorable site to transmigrate, and adhesion strengthening aids this process as the neutrophil must resist detachment under the shear stress of blood flow (9, 48, 57, 58). Interestingly, we found that the involvement of vinculin in regulating neutrophil migration is dependent on the hemodynamic context in which it is assayed. While the migration of vinculin-deficient neutrophils was reduced compared to wild-type neutrophils in a commonly used static motility assay, no impairment in migration under flow was observed in vinculin knockout neutrophils. In the flow chamber assay, there was a preference for neutrophils to crawl in the direction of shear stress rather than opposing it, and both wild-type and vinculin-deficient neutrophils exhibited this directional behavior. There is much evidence for *in vivo* chemokine gradients superseding *in vivo* physical cues such as shear stress to direct neutrophils toward sites of injury (56, 59). However, shear stress is still a quantifiable factor *in vivo* with neutrophils often favoring travel with flow rather than against (57). Addressing the chemotactic persistence of neutrophils under flow after vinculin ablation would be critical as this has been ascribed as a critical role for vinculin, which was not resolved with our experiments that employed immobilized CXCL1 as a chemokinetic stimulus (44). In the absence of forces due to shear stress, random migration was observed, with no significant role for vinculin in the persistence of neutrophil migration based on directness and the linearity of the mean-squared displacement graph (60). However, the model in which mean-squared displacement was calculated has a limited fit for calculations at greater time intervals, and the curvature of mean-squared displacements for wild-type and vinculin knockout neutrophils might not be completely solved.

Outside-in signaling by integrins plays a prominent role in actin cytoskeletal reorganization for physiological functions such as migration. Here we show a marked ablation of stable F-actin and neutrophil polarization on ICAM-1 in vinculin knockout neutrophils that may indicate an impaired outside-in integrin signaling response. Stable F-actin is important for the contractility of neutrophils and force generation is attenuated in vinculin knockout neutrophils as quantified by traction force microscopy. Outside-in signaling by ligand-bound integrins is thought to occur through ITAM-containing receptors, such as the low affinity Fc receptor Fcγ and DAP12 expressed in leukocytes (61). These receptors are phosphorylated by Src family kinases (SFKs) to recruit and activate Syk, leading to downstream activity of PLCγ2, Vav exchange factors, PI3K, and SLP-76 adaptor. In this way SFKs are responsible for adhesion strengthening, morphological change, and migration through outside-in signaling. However, while Syk has been found to localize with proteins such as the β2 integrin Mac-1 (CD11b/CD18) and mAbp1, which are important for intraluminal crawling, the proteins necessary for integrin outside-in signaling (Syk, SFKs and ITAM-containing receptors) have been found to have disparate roles in intraluminal crawling and only minimally influence extravasation in fMLP- and chemokine-mediated models of neutrophil recruitment (62). For example, a recent study found the Tec family kinase Btk and the SFK Hck, while indispensable for fMLP-mediated intraluminal crawling and recruitment, were dispensable for chemokine-mediated neutrophil recruitment (63). Our study is limited to chemokine-mediated neutrophil recruitment, so we cannot rule out other pathways of activation that might involve vinculin-dependent intraluminal crawling or outside-in signaling in general. Interestingly, while PKCθ and mAbp1 are implicated strongly in adhesion strengthening and intraluminal crawling under flow conditions, respectively, they have more dispensable roles under static conditions (48, 57). Here we describe the reverse, in which vinculin has a dispensable role for neutrophil crawling under shear forces but is important for crawling in the absence of flow *in vitro*. Together with these previous findings, our data suggests that neutrophils employ distinct mechanisms for migration that depend upon microenvironmental conditions and cues. One might speculate that vinculin is more important to interstitial crawling compared to vascular recruitment, but additional studies are necessary to directly address this possibility. The *in vitro* system used in our study to probe neutrophil migration is two-dimensional and used ligands to mimic endothelium, while interstitial crawling in tissues is most often three-dimensional and can have a dispensable role for integrins (64).

Outside-in and inside-out signaling have been defined for integrins to describe their nature as both adhesion molecules and signaling receptors (65). We find that vinculin was not required to regulate neutrophil β2 integrin activation through inside-out signaling when neutrophils were activated by soluble CXCL1. We cannot rule out that vinculin may participate in integrin regulation through an outside-in pathway once bound to ICAM-1, in which talin-1 is bound to the β2 integrin tail (7). The outside-in component of integrin signaling was of interest to this study because of vinculin’s known role in mechanotransduction in other cell types (51, 66). As neutrophils can travel from the blood to virtually all parts of the body, sensing the mechanical environment is of immense importance to fine tune neutrophil function in different tissues (31). In mesenchymal cells, rigidity sensing is a well-characterized adaptive response that influences focal adhesion formation, cell spreading, and traction force generation (67). Indeed, our data indicate that vinculin can also mediate mechanosensing by neutrophils. However, the stiff gels of 100 kPa used in our study are not of a physiologically relevant tissue compliance. Vinculin-dependent spreading was only found to be impaired on 100 kPa substrates, while traction stress generation and spreading was not impaired on physiologically relevant substrate stiffnesses between 1-10 kPa. Rescue of vinculin-deficient of neutrophil spreading was achieved through exogenous expression of full-length vinculin, but not by the A50I mutant of vinculin that disrupts its interaction with talin-1 (17). These data point towards a mechanism that involves the function of vinculin at integrin-based adhesion structures. A mechanosensing phenotype was also resolved using traction force microscopy, with a small, but significant, attenuation of neutrophil contraction observed after vinculin depletion. This defect would be unlikely to affect motility, which agrees with the *in vivo* model and suggests that physiological neutrophil motility is unimpaired in the absence of vinculin. Considering that vinculin expression is unnecessary for neutrophil recruitment using a murine model of acute peritonitis, it remains unclear whether this might be because rigidity sensing by neutrophils does not play a role in their recruitment to this specific tissue.

In conclusion, we report that the role of vinculin in neutrophil adhesive function mediated by β2 integrins is highly dependent on the mechanical context of the microenvironment. Analysis of neutrophil spreading and adhesion under static conditions, using assays commonly used in the field, first suggested a prominent role for vinculin. By performing assays of mechanosensitivity and of adhesion and migration under flow, we observed that vinculin is dispensable when experimental conditions more closely mimic physiological conditions. Finally, we found that vinculin is also not required for neutrophil recruitment in an animal model of sterile inflammation. Further studies are necessary to probe other inflammatory contexts and tissue sites to determine whether a vinculin-dependent mode of neutrophil adhesion and motility is employed *in vivo* under different microenvironmental conditions.

## Acknowledgements

We acknowledge Dr. Paul Ekert for the tamoxifen-inducible Hoxb8 plasmid used to produce the progenitor cell lines used in this study. This study was supported by the following awards from the National Institutes of Health: R35GM124911 (CTL), R01AI116629 (JSR and CF), F31DE028745 (HW), and T32HL134625 (BMN and HW). All authors declare no related conflict of interest.

## Author contributions

ZSW designed and performed experiments, analyzed data, and wrote the manuscript. LH, MP, and MH designed, performed and analyzed traction force microscopy experiments, and were supervised by CF. HW performed and analyzed experiments probing neutrophil spreading on polyacrylamide gels and was supervised by JSR. BMN analyzed some of the neutrophil migration data. AW assisted with adhesion assays. CTL designed experiments, supervised the study and wrote the manuscript.

## Supplementary information

**Supplementary Video 1. Neutrophil migration in a flow chamber.** Flow chambers coated with E-selectin, ICAM-1, and CXCL1 were perfused for 30 minutes with a 1:1 mix of WT (CFSE labeled) and VclKO neutrophils at a wall shear stress of 1 dyne/cm^2^. Scale bar = 10 μm.

**Figure S1.**
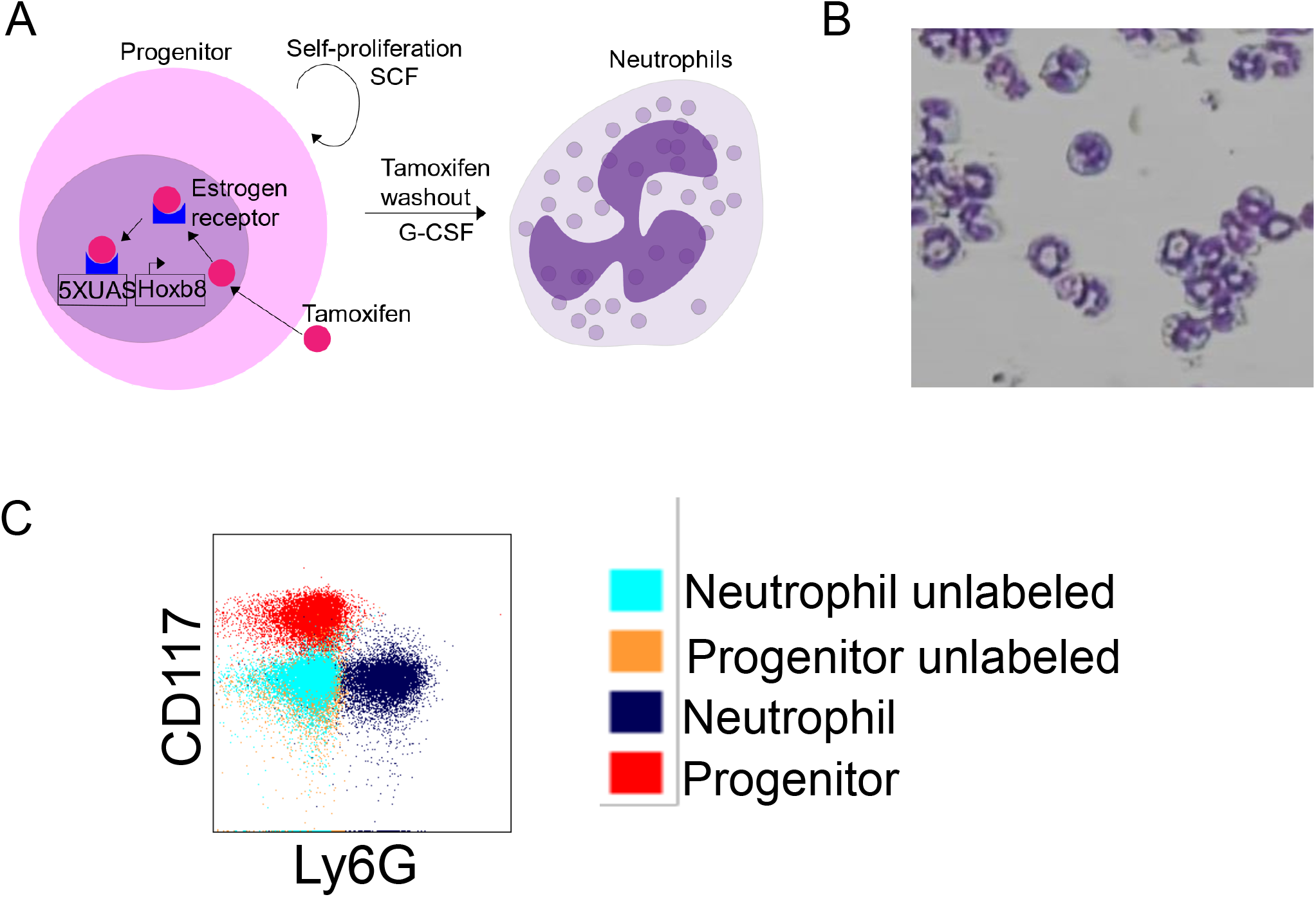
Differentiated HoxB8-conditional progenitors display a neutrophil phenotype. (A) Schematic of the inducible HoxB8 progenitor system and G-CSF-induced differentiation into neutrophils. (B) Wright-Giemsa stain of *in vitro* progenitor-derived neutrophils. (C) Flow cytometry of day 4 differentiated progenitors, analyzing neutrophil marker Ly6G and progenitor marker CD117.

**Figure S2.**
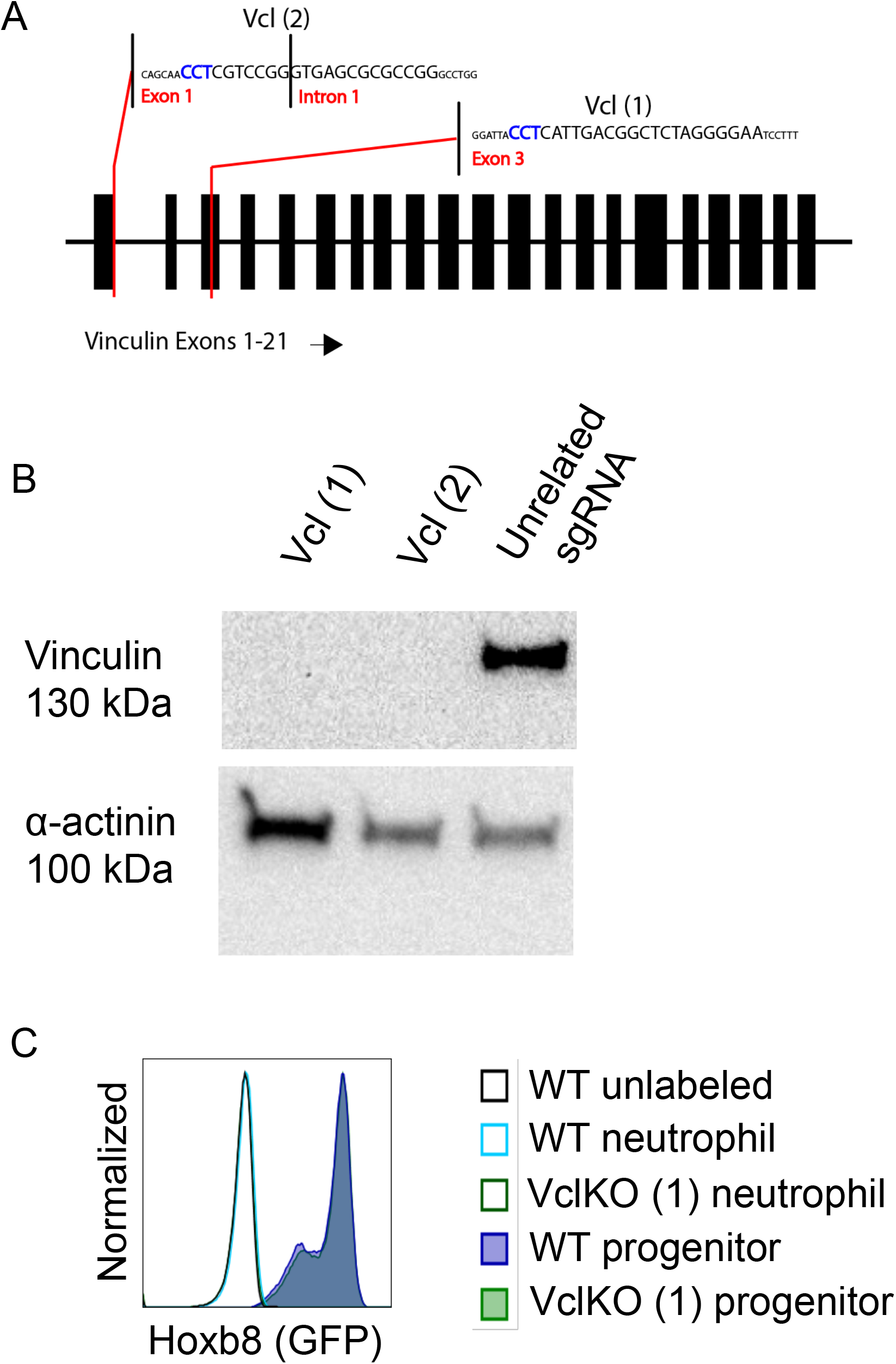
Knockout of vinculin in HoxB8-conditional progenitors and neutrophils. (A) Schematic of CRISPR/Cas9 targeting of the murine Vcl locus with independent sgRNAs. (B) Representative western blot of *in vitro*-derived neutrophil protein expression of vinculin and loading control α-actinin. (C) Representative flow cytometry histograms of wild-type (WT) and vinculin knockout (VclKO) progenitors and neutrophils, analyzing HoxB8-GFP expression (representative of 3 independent experiments).

**Figure S3.**
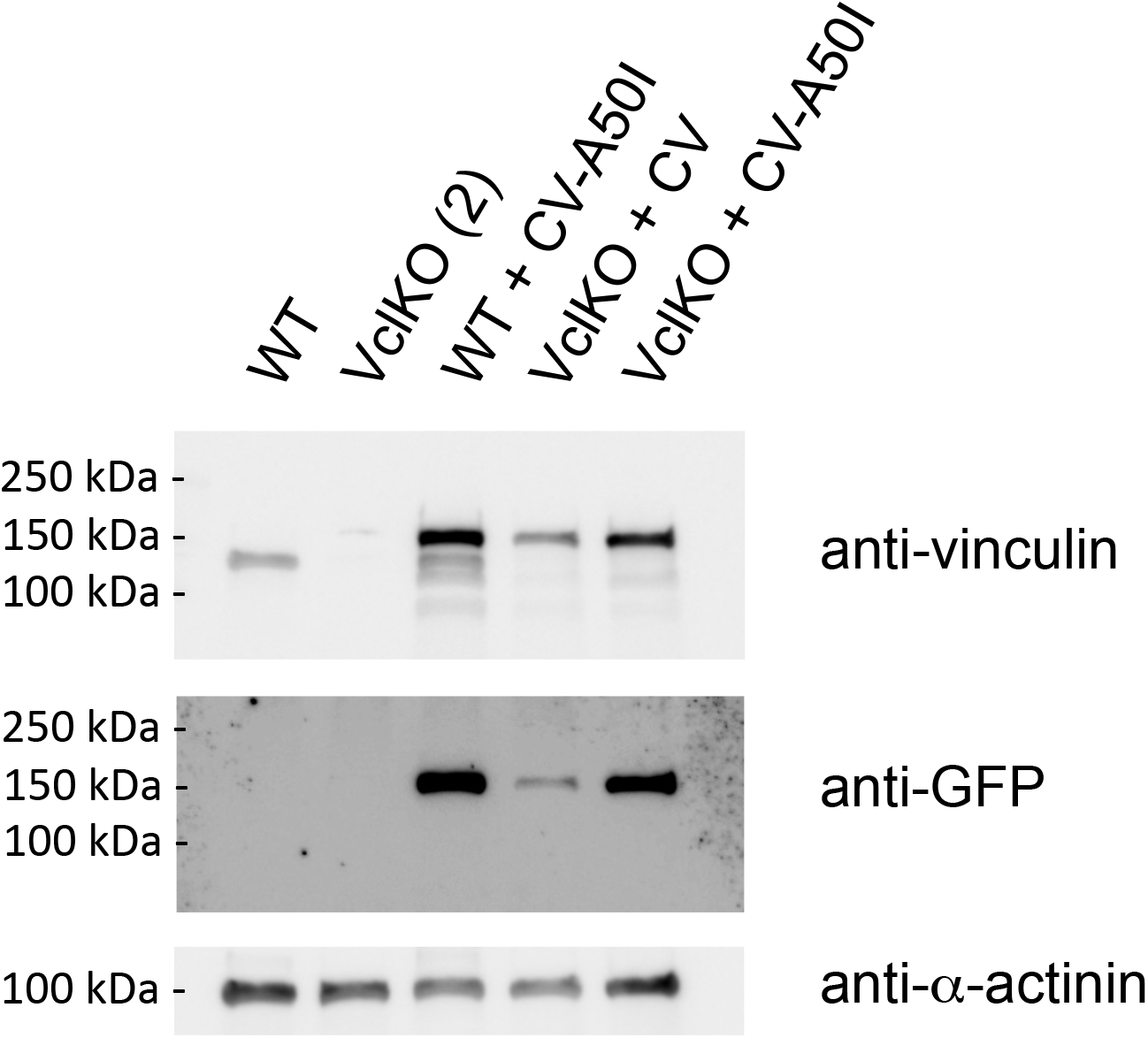
Exogenous re-expression of Clover-vinculin. Representative western blot of *in vitro*-derived neutrophil protein expression of vinculin (endogenous, recognized by anti-vinculin), Clover-vinculin (exogenous, recognized by both anti-vinculin and anti-GFP), and loading control α-actinin. Clover-vinculin is abbreviated CV.

**Figure S4.**
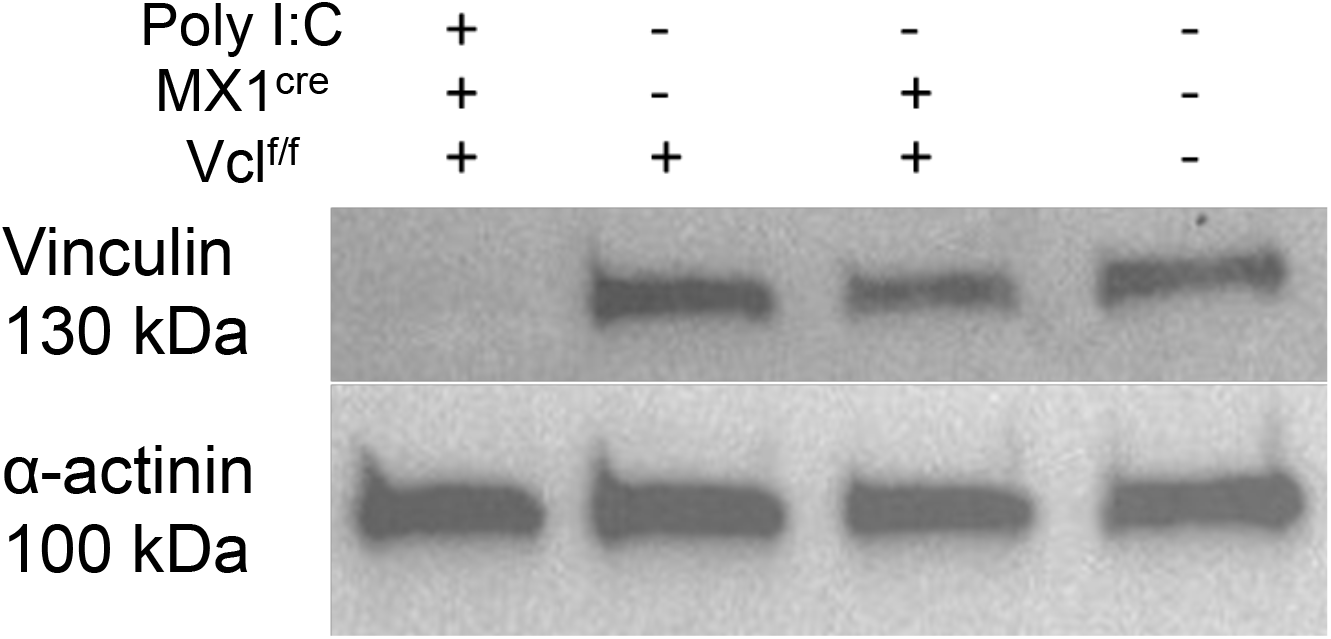
Tissue-specific knockout of vinculin in mice. Representative western blot of purified bone marrow neutrophils from control, Vcl^f/f^, and Vcl^f/f^MX1^cre^ mice, with and without Poly I:C activation of MX1-Cre recombinase-induced disruption of *Vcl*. Western blots were probed for protein expression of vinculin and loading control α-actinin.

**Figure S5.**
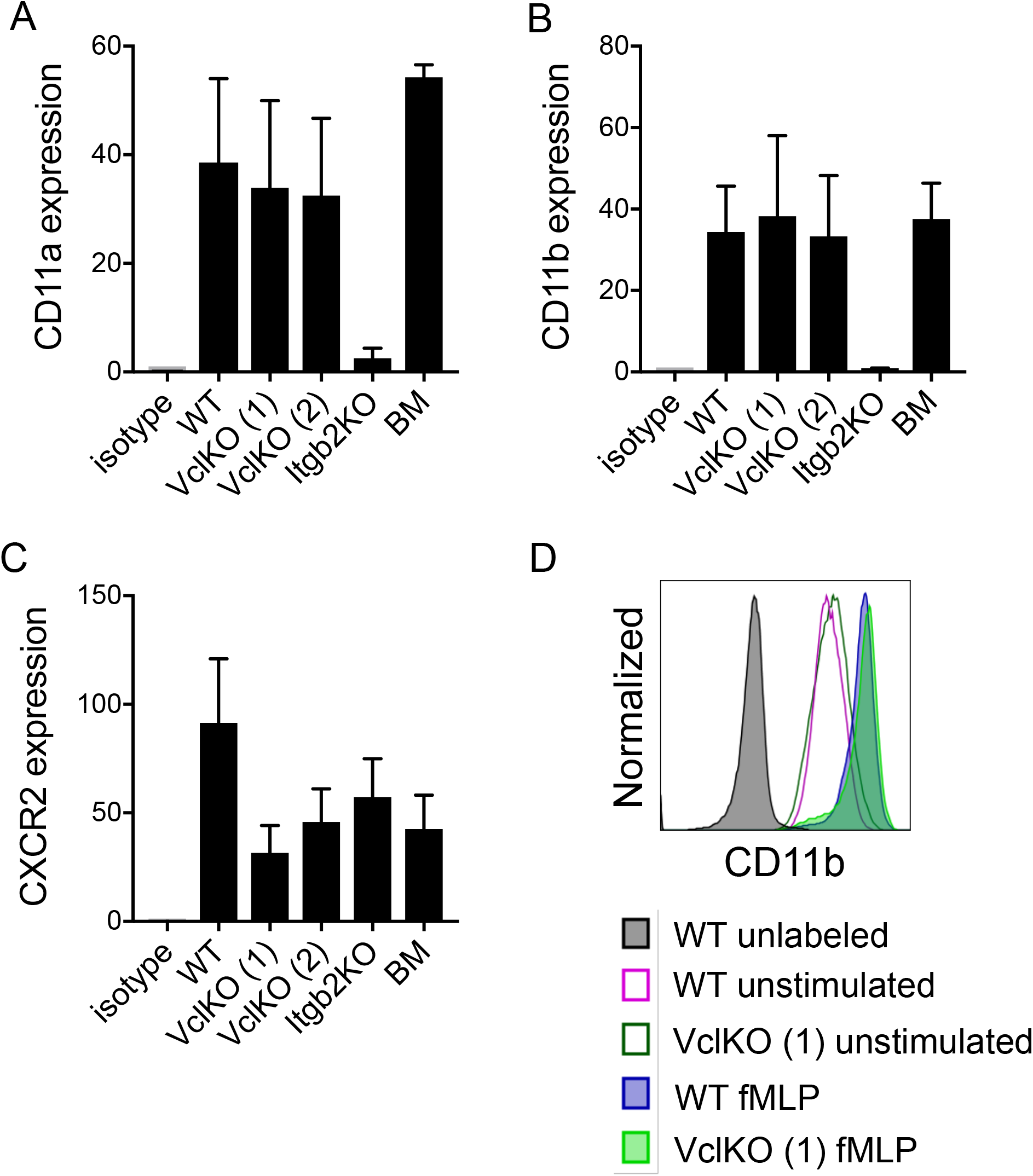
Expression of adhesion and signaling receptors in progenitor-derived neutrophils. Flow cytometry quantification of cell surface expression of (A) CD11a, (B) CD11b, and (C) CXCR2 by mean fluorescent intensity (MFI). Groups of *in vitro*-derived neutrophils are indicated and are shown with analyses of bone marrow neutrophils (n=4 independent experiments, except for BM where n=2). (D) Flow cytometry analysis of CD11b upregulation after fMLP stimulation for WT and VclKO neutrophils (representative of n=2 experiments).

**Figure S6.**
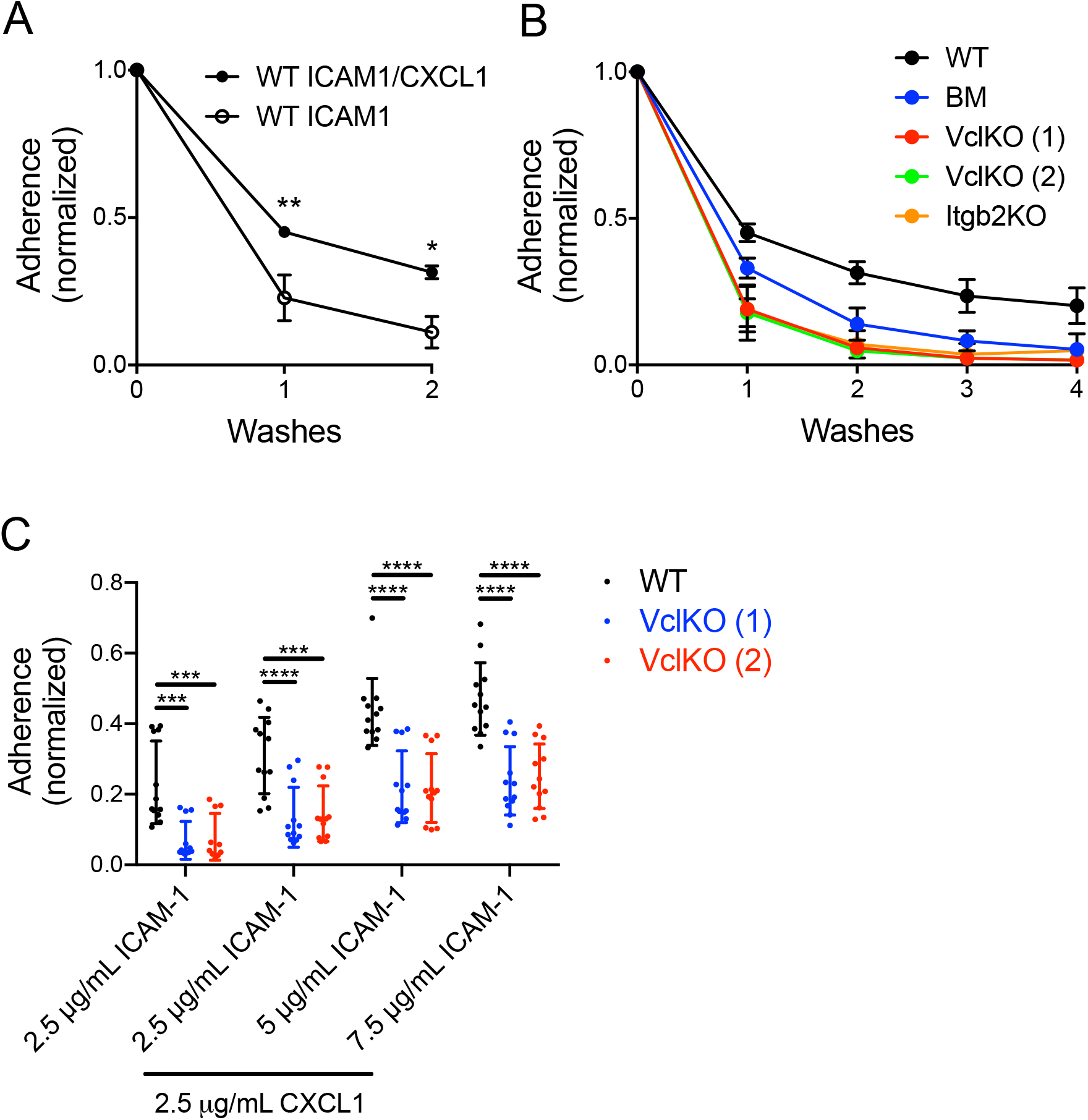
Additional analyses of neutrophil static adhesion assays. (A-C) Normalized adhesion of CFSE-labeled neutrophils under conditions as indicated: (A) Evaluation of CXCL1-induced neutrophil adhesion to ICAM-1, (B) Analysis of sequential washes to remove non-adherent neutrophils, (C) Analysis of coating concentration dependence of neutrophil adhesion. All experiments were analyzed using one-way or two-way ANOVA with Tukey multiple comparison test (3 independent experiments). * p<0.05; ** p<0.01; *** p<0.001; **** p<0.0001.

**Figure S7.**
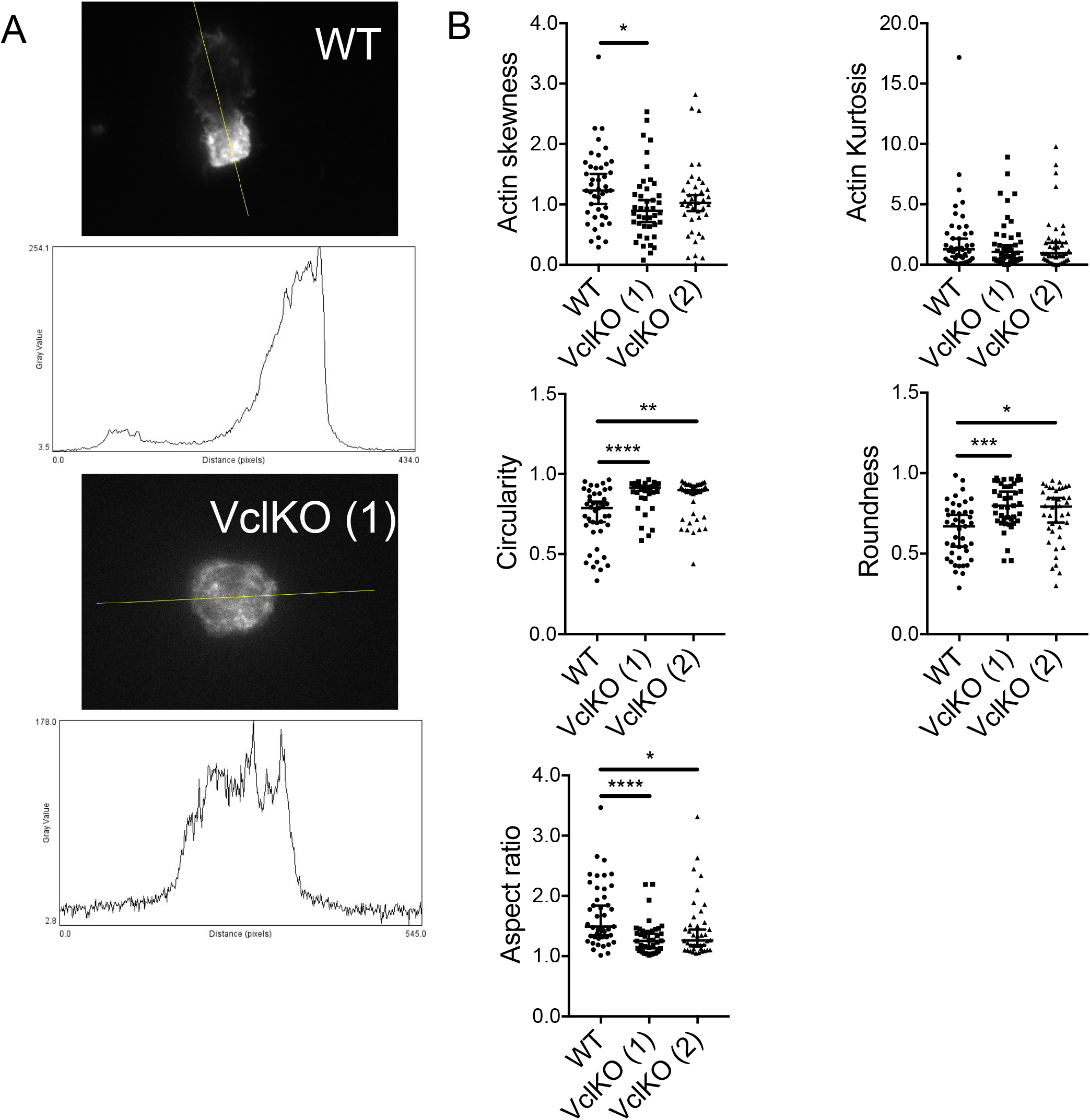
Vinculin-deficient neutrophils display a less polarized phenotype. (A) Representative image of F-actin distribution (by TIRFM) of WT and VclKO neutrophils, displayed with a cross-sectional histogram of fluorescence intensity at the plane indicated. (B) Analyses of the fluorescent distribution of actin and shape of polarized neutrophils based on F-actin staining of neutrophil after 30 minute adhesion and spreading on ICAM-1 and CXCL1. Data were analyzed using Kruskal-Wallis one-way ANOVA on ranks with Dunn’s multiple comparison test (n>30 cells per group, 3 independent experiments). * p<0.05. ** p<0.01. *** p<0.001, **** p<0.0001.

**Figure S8.**
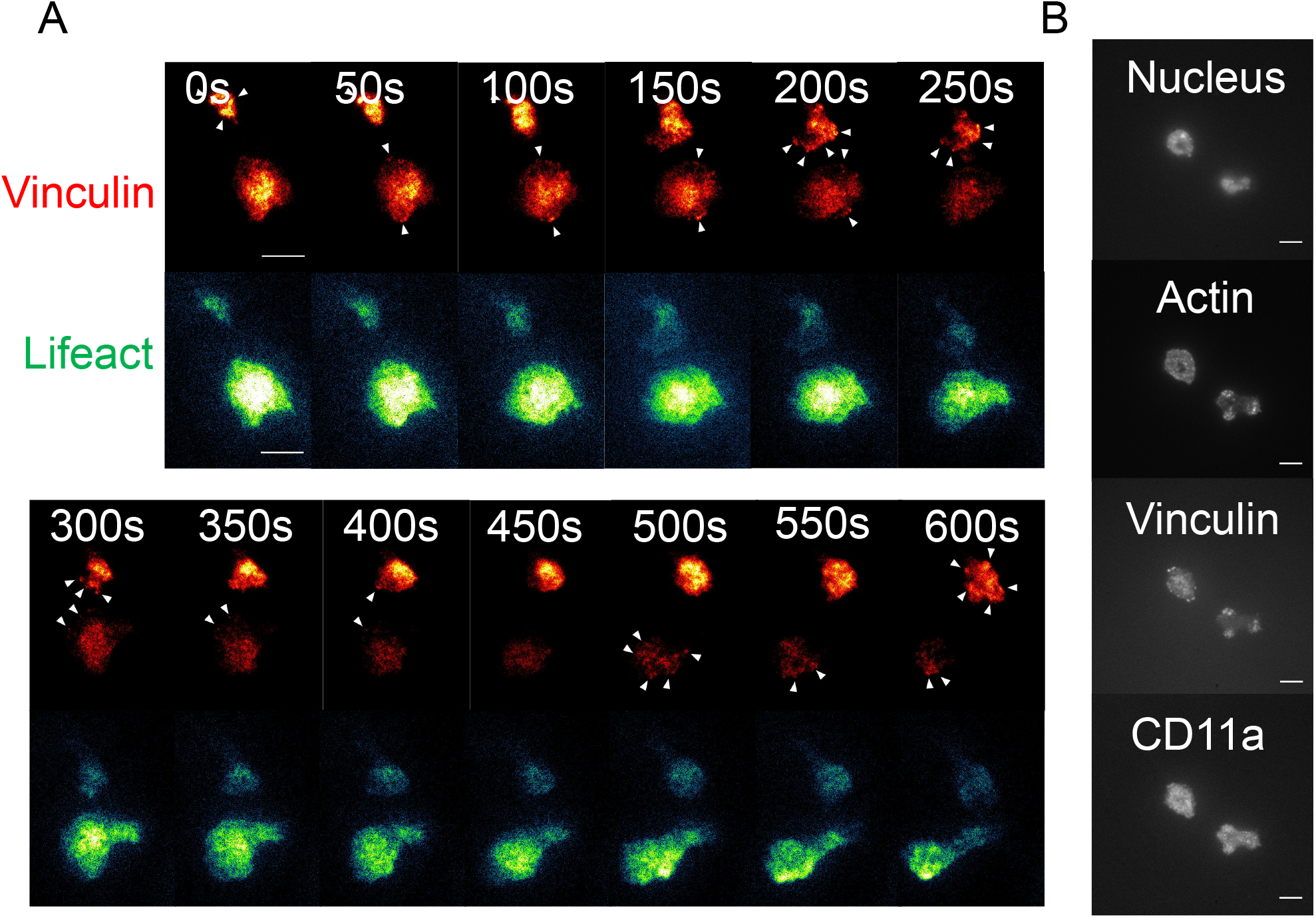
Vinculin localizes to contracting pseudopods. (A) Migration of neutrophils expressing Clover-vinculin and Lifeact-mRuby2 on a substrate of ICAM-1 and CXCL1 over the course of 10 minutes (representative of 3 independent experiments). Arrows point to punctate vinculin-containing structures. Scale bar = 10 μm. (B) Immunocytochemistry of neutrophils fixed after 30 minutes of migration on ICAM-1 and CXCL1. Samples were stained with antibodies against vinculin and CD11a; phalloidin and Hoescht used for F-actin and nuclear staining. Scale bar = 10 μm.

**Figure S9.**
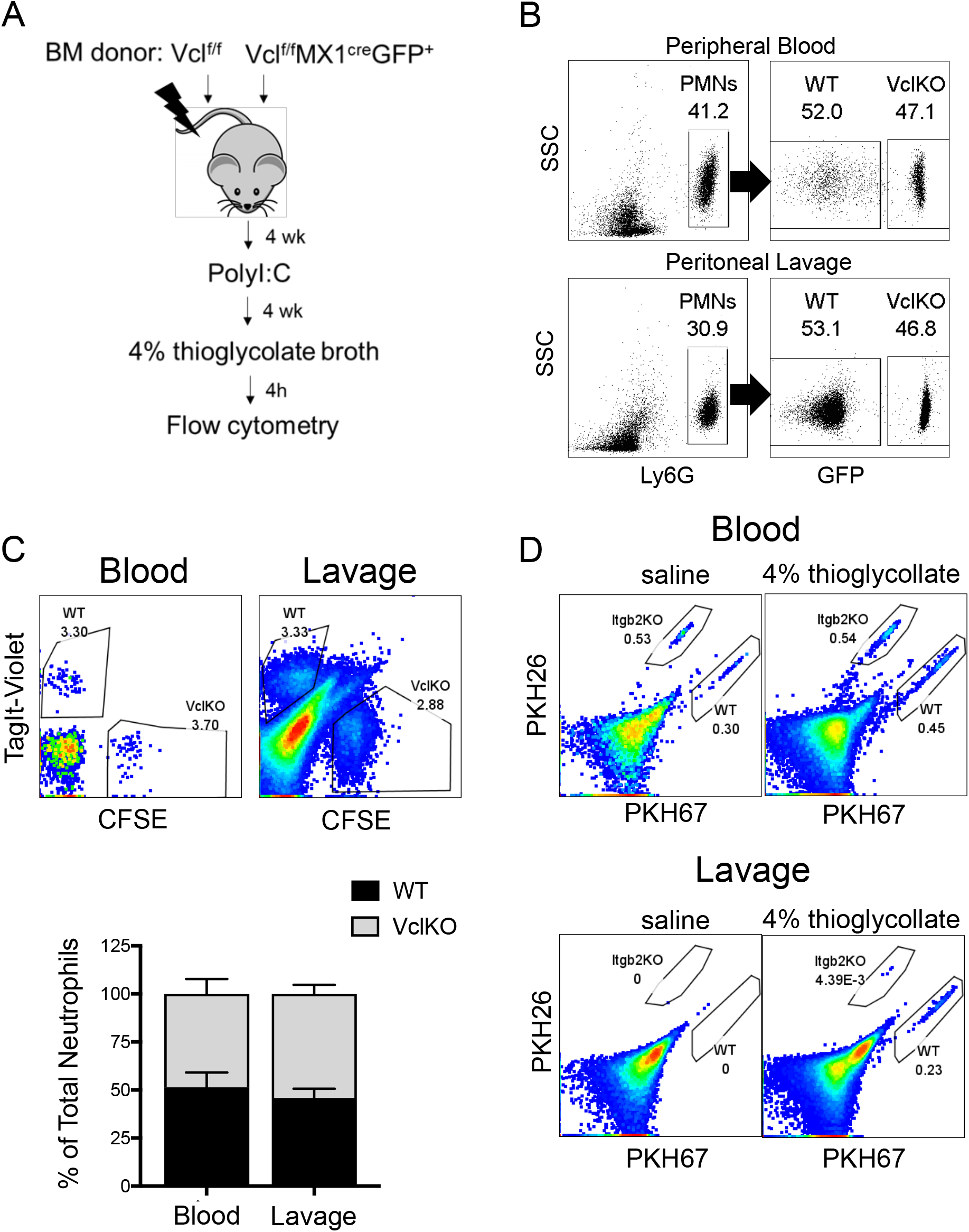
Analyses of neutrophil trafficking during sterile peritonitis. (A) Schematic that depicts the experimental design for analyzing neutrophil trafficking in murine mixed chimeras reconstituted with both control (Vcl^f/f^) and vinculin knockout (Vcl^f/f^MX1^cre^GFP^+^) bone marrow. Induction of vinculin knockout was achieved with Poly I:C treatment. (B) Representative flow cytometry dot plots of peripheral blood and peritoneal lavage of mixed chimeric mice, depicting the method to determine the fraction of wild-type and vinculin knockout neutrophils within those compartments. (C-D) Experiments to determine the trafficking of progenitor-derived neutrophils that were labeled with distinct fluorescent dyes, mixed in 1:1 ratios, and intravenously injected into mice during the course of thioglycollate-induced peritonitis. In (C), representative (top) and composite (bottom) analysis of the frequency of adoptively transplanted TagIt-Violet labeled WT neutrophils (*in vitro*-derived) and CFSE labeled VclKO neutrophils (*in vitro*-derived) in the blood and peritoneal lavage, 4 hours after induction of peritonitis. In (D), representative analysis of the frequency of adoptively transplanted PKH67 labeled WT neutrophils (*in vitro*-derived) and PKH26 labeled Itgb2KO neutrophils (*in vitro*-derived) in the blood and peritoneal lavage, 4 hours after induction of peritonitis. As a control, these same analyses are shown for mice receiving intraperitoneal saline rather than 4% thioglycollate.

**Figure S10.**
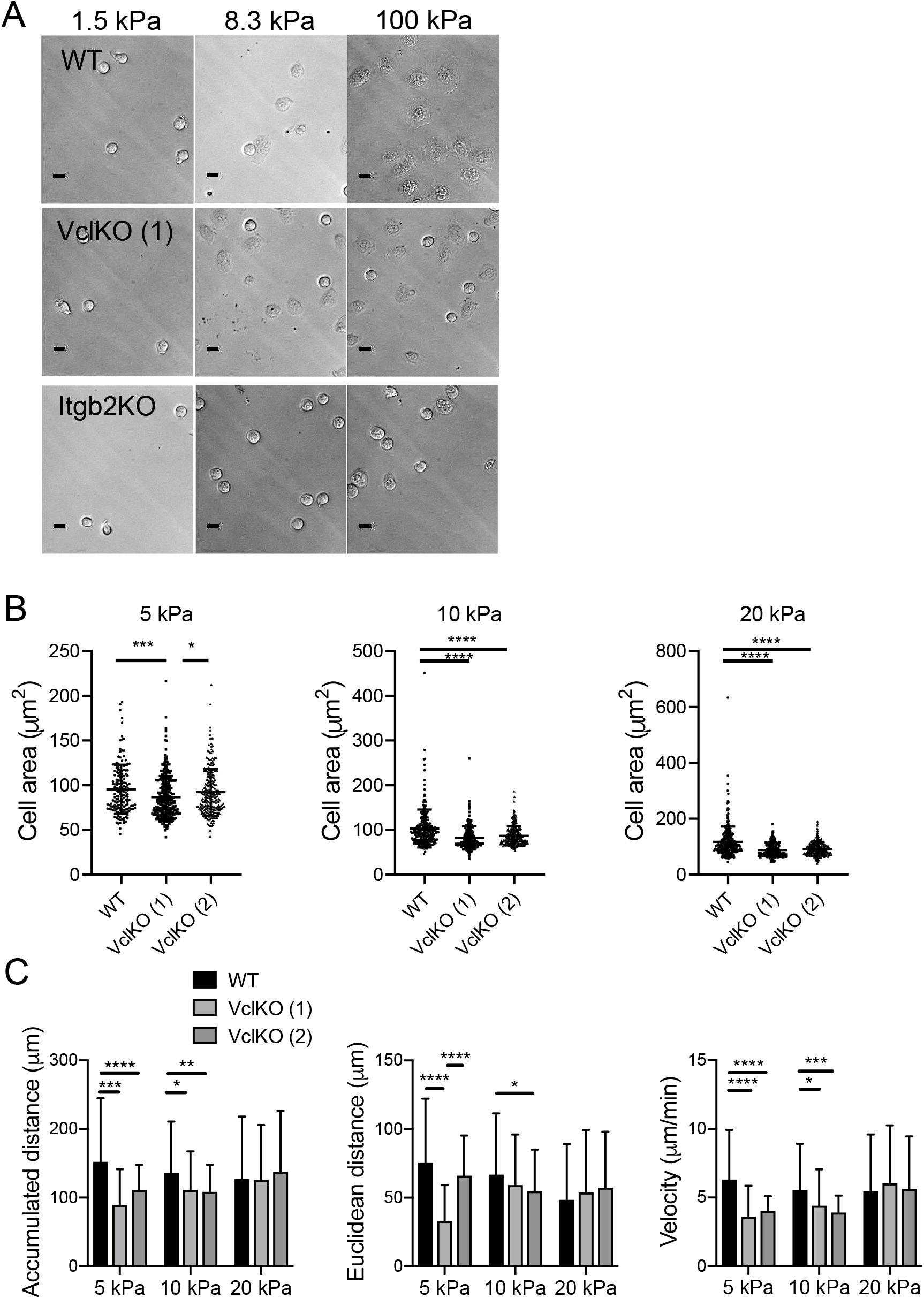
Additional analyses of neutrophil mechanosensing. (A) Representative images of WT, VclKO, and Itgb2KO neutrophil spreading on ICAM-1/CXCL1-conjugated polyacrylamide gels across a range of stiffness, as imaged using a 40X objective, DIC light microscope. Scale bar = 10 µm. (B, C) Further analyses of WT and VclKO neutrophil spread area and motility parameters (accumulated distance, Euclidean distance, velocity) on ICAM-1/CXCL1-conjugated polyacrylamide gels of intermediate stiffnesses: 5 kPa, 10 kPa, and 20 kPa.

